# Telomere-to-telomere sheep genome assembly reveals new variants associated with wool fineness trait

**DOI:** 10.1101/2024.07.21.604451

**Authors:** Ling-Yun Luo, Hui Wu, Li-Ming Zhao, Ya-Hui Zhang, Jia-Hui Huang, Qiu-Yue Liu, Hai-Tao Wang, Dong-Xin Mo, He-Hua EEr, Lian-Quan Zhang, Hai-Liang Chen, Shan-Gang Jia, Wei-Min Wang, Meng-Hua Li

**Author notes:** These authors contributed equally: Ling-Yun Luo, Hui Wu and Li-Ming Zhao.

## Abstract

Ongoing efforts to improve sheep reference genome assemblies still leave many gaps and incomplete regions, resulting in a few common failures and errors in sheep genomic studies. Here, we report a complete, gap-free telomere-to-telomere (T2T) genome of a ram (*T2T-sheep1.0*) with a size of 2.85 Gb, including all autosomes and chromosomes X and Y. It adds 220.05 Mb of previously unresolved regions (PURs) and 754 new genes to the most updated reference assembly, *ARS-UI_Ramb_v3.0*, and contains four types of repeat units (SatI, SatII, SatIII, and CenY) in the centromeric regions. *T2T-sheep1.0* exhibits a base accuracy of >99.999%, corrects several structural errors in previous reference assemblies, and improves structural variant (SV) detection in repetitive sequences. We identified 192,265 SVs, including 16,885 new SVs in the PURs, from the PacBio long-read sequences of 18 global representative sheep. With the whole-genome short-read sequences of 810 wild and domestic sheep representing 158 global populations and seven wild species, the use of *T2T-sheep1.0* as the reference genome has improved population genetic analysis based on ∼133.31 million SNPs and 1,265,266 SVs, including 2,664,979 novel SNPs and 196,471 novel SVs. *T2T-sheep1.0* improves selective tests by detecting several novel genes and variants, including those associated with domestication (e.g., *ABCC4*) and selection for the wool fineness trait (e.g., *FOXQ1*) in tandemly duplicated regions.

## Introduction

Among the first domesticated livestock species, sheep (*Ovis aries*) have evolved various phenotypes, providing an important source of meat, fur, and dairy products^1^. A reference genome assembly of sheep is essential for exploring the evolutionary history^2^, migration^3^, genetic diversity^4^, and causative genes and variants underlying specialized traits^5^ of sheep. With the rapid advancement of sequencing technologies such as high-throughput chromosome conformation capture (Hi-C), Pacific Biosciences (PacBio) and Oxford Nanopore Technology (ONT) sequencing, continuous efforts have been made to improve sheep reference genomes. To date, as many as 57 sheep assemblies at the chromosome or scaffold level have been made available in public databases, including the most updated genomes, such as *Oar_v4.0* (GenBank accession no. GCA_000298735.2)^6^, *Oar_rambouillet_v1.0* (GCF_002742125), and *ARS-UI_Ramb_v2.0* (GCA_016772045.1)^7^.

However, these sheep assemblies suffer from numerous gaps, misassembled regions, uneven sequence depth, varied alignment rates, and mapping failures and errors^4,8^. The total size of the unplaced contigs and scaffolds could be as large as hundreds of million bases. In particular, numerous regions enriched in highly repetitive sequences, such as centromeres, telomeres and transposable elements (TEs), remain unresolved. Additionally, the draft Y chromosome with a size of 25.92 Mb was recently updated^9^ in *ARS-UI_Ramb_v3.0* (GCA_016772045.2, *Ramb_v3.0*), but the divergence and structure of the sheep Y chromosome is still to be confirmed due to abundant repeats, such as long interspersed nuclear elements (LINEs) and long terminal repeats (LTRs)^10,11^.

Through the use of sequencing for ultralong reads and assembly algorithms^12^, telomere-to-telomere (T2T) genome assemblies have been achieved. Accordingly, previously unresolved/unassembled genomic regions, which are enriched in centromeric satellites, nonsatellite segmental duplications and rDNAs^14–16^, as well as novel genes and variants have been revealed in several species, including human^13,14^, ape^15^, maize^16^, Arabidopsis^17^, soybean^18^, and rice^19^. However, a complete gap-free T2T ovine genome has not been available until now.

Here we report the *de novo* T2T gap-free genome assembly for a ram (HU3095) of Hu sheep (*T2T-sheep1.0*), a well-known, highly prolific breed native to China. This complete and seamless assembly covers the Y chromosome and was achieved by using ONT and PacBio HiFi reads of a ram and the sequenced genomes of its parents. Accordingly, the haplotype genome assemblies *T2T-sheep1.0P* and *T2T-sheep1.0M* were also assembled at T2T level. The *T2T-sheep1.0* genome assembled here was subsequently used as a reference to investigate genomic components, particularly centromere and telomere structures, previously unresolved genomic regions, novel genes, structural variants (SVs) and single nucleotide polymorphisms (SNPs). Furthermore, the application of *T2T-sheep1.0* in the analysis of wild and domestic sheep populations worldwide showed its advantages over previous assemblies in variant calling, population genomics analyses and identification of novel genes and variants associated with particular phenotypic traits (e.g., wool fineness) under selection.

## Results

### T2T gap-free genome assembly

A total of 543.2 Gb of ultralong ONT reads (190.4× coverage) and 149.0 Gb of PacBio HiFi reads (52.2× coverage) were obtained to assemble the *T2T-sheep1.0* reference genome (Supplementary Table 1, Supplementary Fig. 1 and Supplementary Methods). Initial assembly was achieved based on the PacBio HiFi data, and consists of 246 contigs with an N50 of 96.54 Mb (Supplementary Table 2). Furthermore, 1135.86 Gb of Bionano’s optical genome mapping (OGM) data and 357.22 Gb of high-throughput chromatin capture (Hi-C) sequencing data (Supplementary Table 1) were used to scaffold the contigs and anchor them onto the 27 pseudomolecules, which correspond to 26 autosomes and chromosome X (ChrX). A total of 139 gaps were identified in the initial assembly, ranging from 108 bp to 1.02 Mb with a total length of 3.41 Mb. The gaps are enriched in centromeric regions that contain highly repetitive sequences (Supplementary Table 3). Ultralong ONT reads were then used to fill the gaps via the strategy of extension or local assembly. Alignments of Bionano optical maps and ONT, HiFi and Hi-C reads indicated that all the gaps in the regions had been filled (Fig. 1a, Supplementary Fig. 2, and Supplementary Fig. 3).

**Fig. 1.**
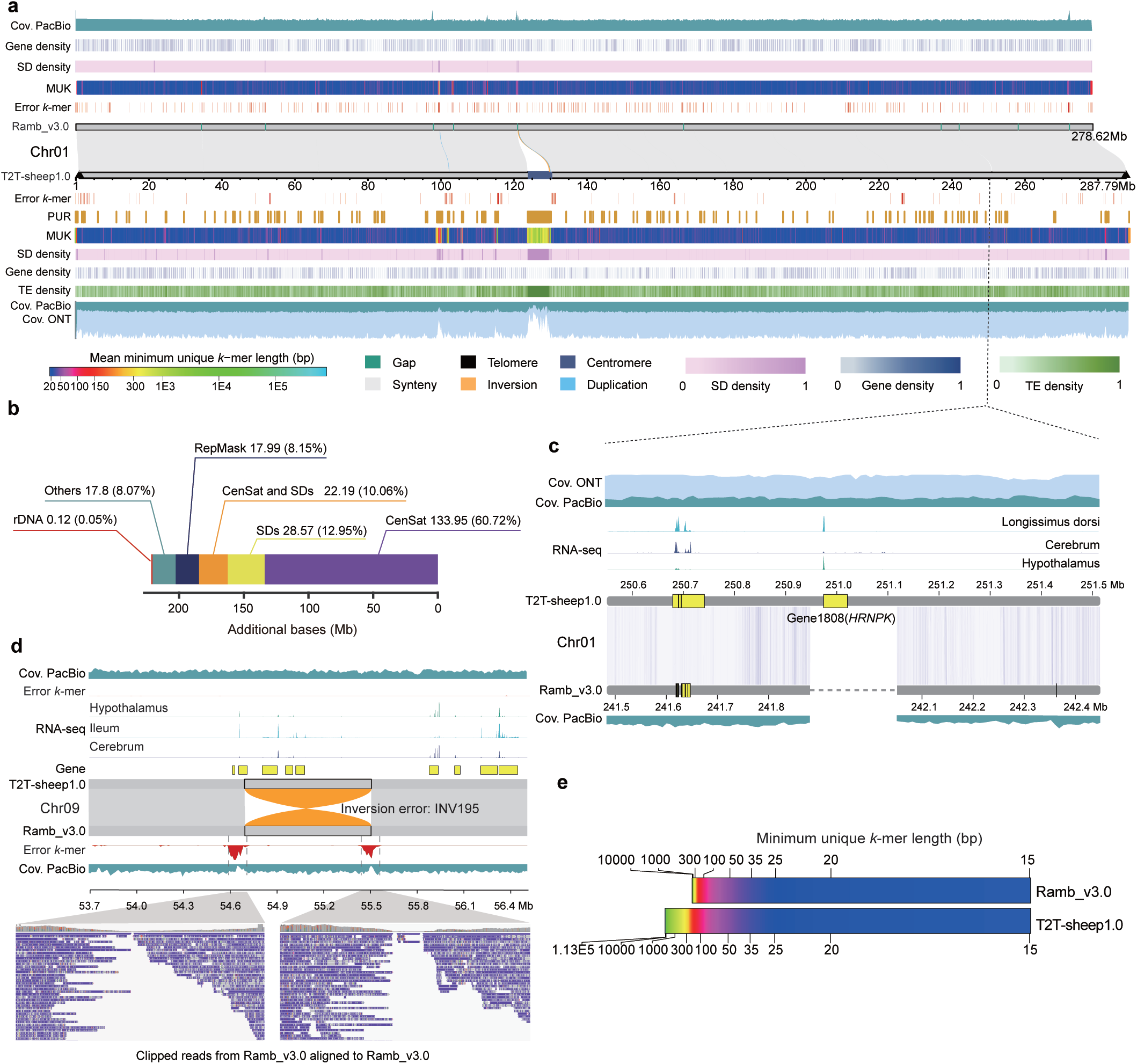
Genomic comparisons of the *Ramb_v3.0* and *T2T-sheep1.0* sheep assemblies. **a,** Genomic features annotated on chromosome 1 (Chr01) for *Ramb_v3.0* and *T2T-sheep1.0*. The coverages of ultralong ONT (Cov. ONT) and PacBio HiFi (Cov. PacBio) long reads are shown in 200-kb windows. Gene, segmental duplication (SD) and transposable element (TE) density values were calculated in 10-kb windows. MUK, minimum unique *k*-mer length in 100-kb windows. Error *k*-mer, *21*-mer errors caused by the worse assembly. PUR, previously unresolved region in *T2T-sheep1.0* compared to *Ramb_v3.0*. The gray blocks between the two horizontal grey bars of *T2T-sheep1.0* and *Ramb_v3.0* indicate collinearity, and one inversion and two duplications between the two assemblies are plotted in orange and blue respectively. Centromeres are highlighted in dark blue, telomeres are marked with black triangles on the “*T2T-sheep1.0*” grey bar, and gaps are shown in green on the *Ramb_v3.0* grey bar. **b,** Contents of various sequence types in the PURs of *T2T-sheep1.0* compared to *Ramb_v3.0*. CenSat, satellites in centromeric regions identified by RepeatMasker. SDs, segmental duplicaitons. RepMask, other repeats identified by RepeatMasker. **c,** One gap containing a gene (Gene1808, namely, *HRNPK*) with transcriptional expression on Chr01 of *Ramb_v3.0* was filled in *T2T-sheep1.0*. Genes colored with yellow showed transcriptional expression according to RNA-seq in 10-kb windows in longissimus dorsi, cerebrum, and hypothalamus tissues. The coverage of ONT and PacBio HiFi reads confirmed the reliability of gap filling. **d,** An inversion error (INV195), highlighted in orange, was found on chromosome 9 (Chr09) in *Ramb_v3.0* and corrected in *T2T-sheep1.0*. The genes in the region were expressed in hypothalamus, ileum, and cerebrum tissues. Two peaks of error *k*-mers (*k* = 21) were found in *Ramb_v3.0* and corresponded to the two junction sites of this false-positive inversion, which cannot be covered by PacBio reads from Rambouillet sheep (NCBI Biosample ID SAMN17575729) assembled previously for *Ramb_v3.0*. **e,** Genome-wide MUK lengths in a comparison of *T2T-sheep1.0* and *Ramb_v3.0*.

Initial assembly of chromosome Y (ChrY) was performed independently based on the paternal-specific ultralong ONT reads. Accordingly, gaps were filled using the Y-chromosome-specific contigs, which were assembled by the trio-binning model of Hifiasm^20^ (v0.14) based on the HiFi long reads (Supplementary Table 1). All 56 telomeric regions were locally assembled based on the HiFi reads, and all the incomplete chromosomal ends were replaced with 56 complete telomeres of 1.20 – 25.32 kb (Fig. 1a and Supplementary Fig. 4). Finally, the complete sheep genome assembly, *T2T-sheep1.0*, with a size of 2.85 Gb, was constructed, covering all the autosomes and two sex chromosomes, X and Y (Table 1). In addition, the trio-based assembly was performed to obtain the haplotype-resolved autosomes for *T2T-sheep1.0P* of paternal origin and *T2T-sheep1.0M* of maternal origin. Parent-specific *k*-mers were generated based on parental short reads to bin the ONT and HiFi reads of paternal or maternal origins, which were used to fill 20 gaps for *T2T-sheep1.0P* and 15 gaps for *T2T-sheep1.0M*. Finally, the complete chromosomes X and Y in *T2T-sheep1.0* were included in *T2T-sheep1.0M* and *T2T-sheep1.0P* respectively (Table 1).

**Table 1.**
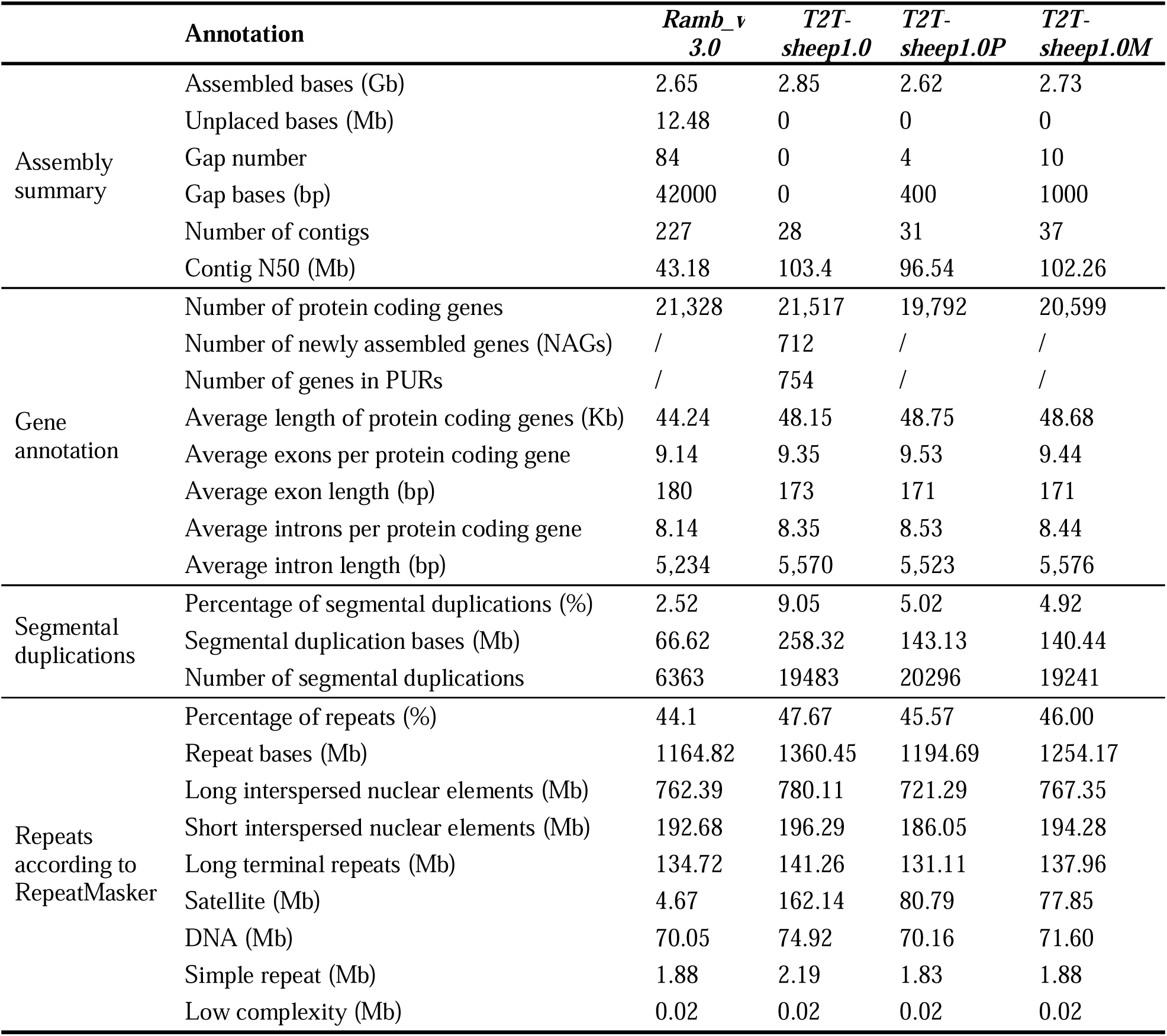
Comparison between Ramb_v3.0, T2T-sheep1.0, T2T-sheep1.0P and T2T-sheep1.0M.

|  | **Annotation** | ***Ramb_v***<br>***3.0*** | ***T2T-***<br>***sheep1.0*** | ***T2T-***<br>***sheep1.0P*** | ***T2T-***<br>***sheep1.0M*** |
| --- | --- | --- | --- | --- | --- |
| Assembly<br>summary | Assembled bases (Gb) | 2.65 | 2.85 | 2.62 | 2.73 |
|  | Unplaced bases (Mb) | 12.48 | 0 | 0 | 0 |
|  | Gap number | 84 | 0 | 4 | 10 |
|  | Gap bases (bp) | 42000 | 0 | 400 | 1000 |
|  | Number of contigs | 227 | 28 | 31 | 37 |
|  | Contig N50 (Mb) | 43.18 | 103.4 | 96.54 | 102.26 |
| Gene<br>annotation | Number of protein coding genes | 21,328 | 21,517 | 19,792 | 20,599 |
|  | Number of newly assembled genes (NAGs) | / | 712 | / | / |
|  | Number of genes in PURs | / | 754 | / | / |
|  | Average length of protein coding genes (Kb) | 44.24 | 48.15 | 48.75 | 48.68 |
|  | Average exons per protein coding gene | 9.14 | 9.35 | 9.53 | 9.44 |
|  | Average exon length (bp) | 180 | 173 | 171 | 171 |
|  | Average introns per protein coding gene | 8.14 | 8.35 | 8.53 | 8.44 |
|  | Average intron length (bp) | 5,234 | 5,570 | 5,523 | 5,576 |
| Segmental<br>duplications | Percentage of segmental duplications (%) | 2.52 | 9.05 | 5.02 | 4.92 |
|  | Segmental duplication bases (Mb) | 66.62 | 258.32 | 143.13 | 140.44 |
|  | Number of segmental duplications | 6363 | 19483 | 20296 | 19241 |
| Repeats<br>according to<br>RepeatMasker | Percentage of repeats (%) | 44.1 | 47.67 | 45.57 | 46.00 |
|  | Repeat bases (Mb) | 1164.82 | 1360.45 | 1194.69 | 1254.17 |
|  | Long interspersed nuclear elements (Mb) | 762.39 | 780.11 | 721.29 | 767.35 |
|  | Short interspersed nuclear elements (Mb) | 192.68 | 196.29 | 186.05 | 194.28 |
|  | Long terminal repeats (Mb) | 134.72 | 141.26 | 131.11 | 137.96 |
|  | Satellite (Mb) | 4.67 | 162.14 | 80.79 | 77.85 |
|  | DNA (Mb) | 70.05 | 74.92 | 70.16 | 71.60 |
|  | Simple repeat (Mb) | 1.88 | 2.19 | 1.83 | 1.88 |
|  | Low complexity (Mb) | 0.02 | 0.02 | 0.02 | 0.02 |

After polishing *T2T-sheep1.0* with ONT and HiFi long reads and NGS short reads, the consensus base quality value (QV) across the whole genome is 51.53, with a QV range of 46.05 to 59.75 for the chromosomes (Supplementary Table 4). An accuracy of > 99.999% for each base and a completeness of 92.761% for the whole assembly were obtained. On average, we obtained 99.75%, 98.99%, and 99.97% mapping rates for short, ONT, and HiFi reads, respectively, against *T2T-sheep1.0*. The even coverage distributions of ONT and PacBio HiFi reads suggest a reliable and continuous assembly (Fig. 1a and Supplementary Fig. 2), and both the Hi-C and Bionano optical map data show high consistency of the overall alignment against the pseudochromosomes in *T2T-sheep1.0* (Supplementary Fig. 3). Together, all the statistics above indicated the reliability and completeness of *T2T-sheep1.0* assembled here. Meanwhile, *T2T-sheep1.0P* and *T2T-sheep1.0M* achieve the T2T level, only with 4 and 10 gaps left in the centromeric or pericentromeric regions of 2 (Chr25 and Chr26) and 6 (Chr10, Chr13, Chr17, Chr23, Chr25, and Chr26) chromosomes respectively due to the lack of binned long reads. The average genome-wide QVs of *T2T-sheep1.0P* and *T2T-sheep1.0M* are 55.41 and 55.23 respectively (Supplementary Table 4). We calculated the switching errors and obtained the estimates of 0.3781‰, 0.0895‰, and 0.1199‰ for *T2T-sheep1.0*, *T2T-sheep1.0P*, and *T2T-sheep1.0M*, respectively. Heterozygous regions between *T2T-sheep1.0P* and *T2T-sheep1.0M* were observed (Supplementary Fig. 5a), as 9,982,198 single nucleotide variants (SNVs), 1,248,272 small insertions and deletions (< 50 bp) and 20155 SVs (≥ 50 bp) were discovered between these two haplotype genomes. The coverage distributions of ONT and PacBio HiFi reads suggest a good quality of two parental genome assemblies, with a few of the potential issues in *T2T-sheep1.0P* and *T2T-sheep1.0M* (Supplementary Fig. 5b). In summary, *T2T-sheep1.0* represents a better assembly for the downstream analysis by merging the chromosomal regions from either of the two haplotypes.

### Improvement of *T2T-sheep1.0*

Good collinearity in the syntenic regions was observed between *T2T-sheep1.0* and the NCBI sheep genome reference *Ramb_v3.0*. Nearly 220.05 Mb of previously unresolved regions (PURs) were identified on all 28 chromosomes of *T2T-sheep1.0*, which were unassembled (i.e., gaps) or misassembled on the chromosomes of *Ramb_v3.0* (Fig. 1a and Supplementary Fig. 2). These PURs are mostly located in the centromeric regions and regions enriched for repeats, including unfinished chromosomal ends of telomeric regions and 81 gaps in *Ramb_v3.0* (Supplementary Fig. 6). Chr26 and Chr15 showed the longest accumulated PURs of 22.57 Mb and 21.41 Mb, respectively, and 5.82 Mb of PURs were identified on ChrY. We did not observe an association between the lengths of the PURs and chromosomes.

The PURs include centromeric satellites (CenSat, 60.72%), segmental duplications (SDs, 12.95%), overlapping CenSat and SDs (10.06%), and other repeats (8.15%) (Fig. 1b). The gaps in *Ramb_v3.0* enriched with repeats were filled in *T2T-sheep1.0*, and some genes were annotated in gap-filled regions of *T2T-sheep1.0* (Fig. 1c and Supplementary Fig. 7). For example, the gene ID “Gene1808” (annotated as *HNRNPK*), located in a gap on Chr01, was expressed in longissimus dorsi, cerebrum, and hypothalamus tissues (Fig. 1c). Overall, hundreds of thousands of gaps and unplaced contigs were observed in the available sheep assemblies (Supplementary Figs 8a and 8b). Compared with these previous assemblies, *T2T-sheep1.0* is not the longest due to sequence redundancy when considering all the chromosomes and unplaced contigs. However, *T2T-sheep1.0* represents the longest complete gap-free sheep genome assembly after removing unknown nucleotides in the gaps and mitochondrial genome sequences (Supplementary Table 5). More than 96% Benchmarking Universal Single-Copy Orthologs (BUSCO) were annotated in *T2T-sheep1.0* based on a total of 9226 core genes in the mammalia_odb10 database, while only 93.9% BUSCO annotation was achieved in *Ramb_v3.0*^7^ and 91.2–92.4% in 15 other sheep genome assemblies (Supplementary Fig. 8c). We compared the total lengths of two known centromeric satellites (GenBank accessions KM272303.1 and U24091) among these assemblies, and *T2T-sheep1.0* has the longest lengths for the two satellites, making it the best assembly in terms of centromeric regions (Supplementary Fig. 8d).

*T2T-sheep1.0* corrected many structural errors in *Ramb_v3.0* (Fig. 1d and Supplementary Fig. 9). Based on the MGI-sequenced short reads, rare *k*-mer errors (*k* = 21) were detected in *T2T-sheep1.0*, while enriched *k*-mer errors indicated potential assembly issues, which could be ascribed to genome-wide structural errors in *Ramb_v3.0*. A few large inversions between the two assemblies were confirmed as structural errors with the support of multiple lines of evidence. For example, the inversion INV195 between *T2T-sheep1.0* and *Ramb_3.0* was detected on Chr09 (Fig. 1d). We retrieved the raw PacBio sequencing data of Rambouillet sheep assembled previously and aligned the reads to *Ramb_v3.0*. The alignment showed clipped reads, and the junction sites of the inversion INV195 in *Ramb_v3.0* could not be covered by the PacBio reads, together with low read coverage and *k*-mer error peaks (Fig. 1d). However, alignment of the PacBio sequences demonstrated a good coverage at the junction sites when against *T2T-sheep1.0* (Supplementary Figs 9a and 9b). Similarly, the alignment of raw HiFi long reads generated from the Hu sheep assembled here further evidence that the region was assembled correctly in *T2T-sheep1.0*, and an inversion error occurred in *Ramb_v3.0*, rather than a breed difference.

The improvement in genome quality is also reflected in the average genome-wide QV of 51.53 in *T2T-sheep1.0*, compared to 44.77 in *Ramb_v3.0* excluding chromosome Y (i.e., *Ramb_v2.0*)^7^. Minimum unique *k*-mers (MUKs) are defined as *k*-mers that occur only once in *T2T-sheep1.0*. Compared to *Ramb_v3.0*, *T2T-sheep1.0* exhibits an overall increase in the number of MUK sequences across the chromosomes (Fig. 1e and Supplementary Fig. 10), e.g., from 155.86 Mb to 156.13 Mb based on *20*-mers, from 234.75 Mb to 236.10 Mb based on *50*-mers and from 263.53 Mb to 277.60 Mb based on *1000*-mers, which might benefit from improvements in both base-level accuracy and the assembly of repetitive regions in *T2T-sheep1.0*. The centromeric and repetitive regions, e.g., SDs, exhibit longer MUKs in 100-kb windows than do the other chromosomal regions (Fig. 1a).

### Genome annotation

Based on the repeat libraries built by combining *de novo* prediction and available repeats in animals, 47.67% (1360.45 Mb) of the *T2T-sheep1.0* genome sequences were identified as repetitive sequences, more than observed (44.10%, 1164.82 Mb) in *Ramb_v3.0* (Table 1). Among the repetitive sequences, a great majority are transposable elements (TEs), with a total size of 1200.48 Mb, accounting for 42.07% of the whole *T2T-sheep1.0* genome. Long interspersed nuclear repeats (LINEs) derived from TEs are the most abundant, spanning 780.1 Mb and covering 27.34% of *T2T-sheep1.0*. The PURs could have been driven by repetitive sequences, which would pose a great challenge for accurate assembly. We found that the repeat contents in PURs were much higher than in the other chromosomal regions (Supplementary Table 6). We assembled satellites with a total length of 162.14 Mb, covering 5.68% of the whole genome. However, a total length of only 4.67 Mb (0.18% of the genome) was found in *Ramb_v3.0*, which is congruent with the high content of satellites in PURs. Centromere-specific satellites and SDs account for 70.78% of the PURs with a predominant distribution (Fig. 1b). Among the PURs of *T2T-sheep1.0*, 286,173 satellite sites (157.10 Mb) and 10,214 SDs (50.76 Mb) account for 96.89% and 19.65%, respectively, of all the satellites and SDs in the whole genome (Supplementary Table 6).

A combination of *ab initio*, homolog-based, and transcriptome-based predictions was used to annotate and integrate the nonredundant gene structures in *T2T-sheep1.0*. After removing the transposon genes, a total of 21,517 high-confidence protein-coding genes were obtained (Table 1), of which 754 are newly anchored genes located in the PURs (Supplementary Fig. 11), and 99% of the protein-coding genes were annotated based on public databases, including the NCBI nonredundant (NR) protein database and Kyoto Encyclopedia of Genes and Genomes (KEGG) database. We searched for newly assembled regions (NARs) on the chromosomes in *T2T-sheep1.0* which are not included in *Ramb_v3.0*^7^ (alignment between all chromosomes of *T2T-sheep1.0* and all chromosomes and unplaced contigs of *Ramb_v3.0*), rather than pairwise chromosomal comparisons for PURs (alignment of all chromosomes between *T2T-sheep1.0* and *Ramb_v3.0*). A total of 712 newly assembled genes (NAGs) were identified among the NARs of *T2T-sheep1.0* (Supplementary Fig. 11). The genes in the PURs and NAGs exhibited transcriptional expression in a diverse set of tissues, e.g., adipose, blood, rumen and hypothalamus based on RNA-seq data (Supplementary Table 7 and Supplementary Fig. 11). We annotated 147 genes within the centromeric regions of 25 chromosomes in *T2T-sheep1.0*, and RNA-seq analysis revealed low expression levels of these genes (Supplementary Table 8).

### Gene families and SDs

For the 18084 orthogroups inferred by the program OrthoFinder (v2.5.5)^21^, we observed obvious expansion of gene families with increased copy numbers in *T2T-sheep1.0* compared to the three genome assemblies of sheep *Ramb_v3.0*, argali *CAU_O.ammon polii_1.0* (*Ovis ammon polii*, GenBank accession no. GCA_028583565.1) and goat *ARS1* (*Capra hircus*, GCF_001704415.1) (Supplementary Table 9). For example, the orthogroup OG0000002 contained 33 genes, with only 1 found in *Ramb_v3.0*, 22 in argali *CAU_O.ammon polii_1.0* and none in goat *ARS1*. Furthermore, gene family expansion showed a strong association with the enrichment of SDs (Supplementary Fig. 12).

To characterize the SD content, we identified 111.06 Mb and 20.55 Mb of nonredundant segmental duplicated sequences on the 28 chromosomes (26 autosomes and chromosomes X and Y) of *T2T-sheep1.0* and *Ramb_v3.0*, respectively. A total of 45.56% of the interchromosomal SDs are scattered across different chromosomes, while 54.44% of the intrachromosomal SDs were identified. The SDs spanned 68.55 Mb, and covered 2622 genes. Among these genes, 44.28% are in paralogous gene groups with more than one gene in *T2T-sheep1.0*, indicating a significant contribution of SDs to gene copy number. Compared with those in *T2T-sheep1.0*, the reductions in SDs and paralogous gene groups in *Ramb_v3.0* might be due to the assembly collapse of repetitive sequences. Of the total SDs, 45.71% overlapped with the PURs of *T2T-sheep1.0*, spanning 50.76 Mb. In particular, 4.52 Mb of SDs within PURs on the Y chromosome were shown to be related to the tandem duplication of three gene families, the testis-specific protein Y-encoded (*TSPY)* gene, the heat shock transcription factor Y-linked (*HSFY*) gene, and the zinc finger Y-linked (*ZFY*) gene. Additionally, our selective sweep tests of global wild and domestic sheep identified strong signals linked to tandemly duplicated genes in the SD-enriched regions (see the results below).

### Centromeric regions and their repeat content

Eleven acrocentric chromosomes shared similar satellite sequences, and created an assembly graph with tangles among them (Fig. 2a), which were further discovered to be centromeric repeats. After gap-filling, the centromeric regions were resolved with the support of sufficient read coverage (Fig. 1a and Supplementary Fig. 2). As histone H3 binds to centromeres on nucleosomes, ChIP-seq based on an antibody of phosphor-CENP-A (Ser7) was used to determine the centromeric regions (Fig. 2b). It is known that centromeres are dominated by centromeric satellites and rich in highly hypermethylated CpG as observed in humans^14,22^. Our evidence also supported the identification of centromeric regions, including the enrichment of methylated cytosine based on HiFi data (Supplementary Fig. 13) and the successful alignment of two known centromere-specific satellite DNA sequences (NCBI accessions KM272303.1 and U24091) in these regions. Hypermethylated regions covered the entire centromeric region on Chr02 (Fig. 2b), and similar to that in humans, the hypomethylation displayed a centromeric dip region (CDR) corresponding to SatII on Chr02. CDR is also reported in humans^22,23^ and medaka fish^24^, as CDRs related to hypomethylation might be associated with interaction with CENP-A for binding of functional kinetochores and serving as a part of distinct marks for euchromatin and heterochromatin^25–27^. Based on the high content of repetitive sequences and their higher similarities in centromeric regions, we performed calculations of pairwise sequence identity and sequence complexity or entropy across all the whole chromosomes. Heatmaps of pairwise sequence identity also confirm centromeric locations with significant continuous blocks (Fig. 2b), while the sequence complexity signals or entropies exhibit different distributions in centromeric regions from those in the other regions (Fig. 2b and Supplementary Fig. 14). These results are congruent with the lines of evidence from enriched signals of ChIP-seq, hypermethylation, and satellite DNA alignment (Supplementary Fig. 13). The centromere lengths ranged from 0.36 Mb to 22.63 Mb, showing no association with chromosomal length (Fig. 2c).

**Fig. 2.**
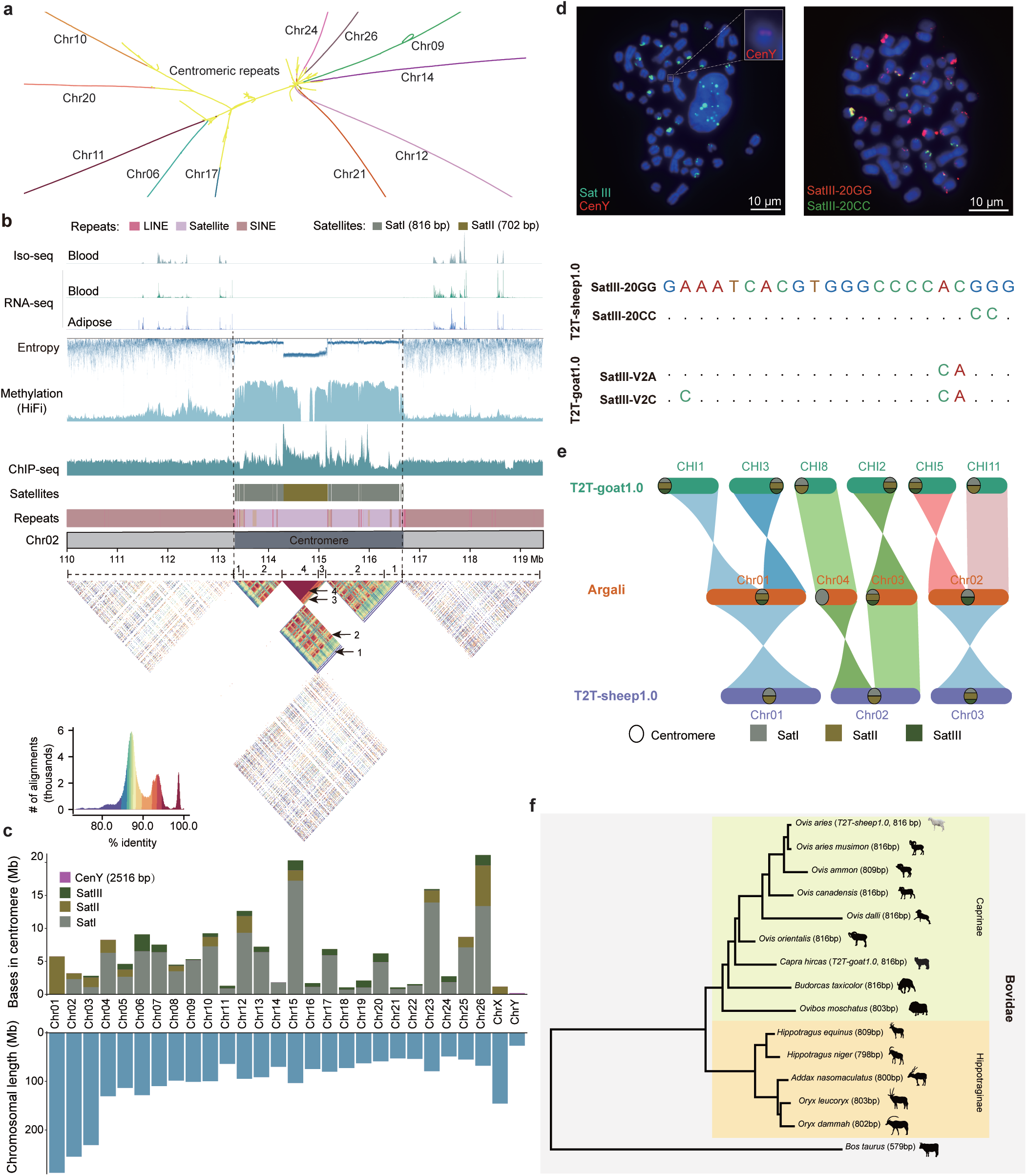
Assembly of centromeric regions and identification of centromeric repeat units. **a,** An assembly graph tangle among the centromeres of 11 acrocentric chromosomes in different colors. Centromeric regions are highlighted in yellow. **b,** Genomic features of the centromeric region on chromosome 2 (Chr02). ChIP-seq for histone H3 variant CENP-A (phospho-CENP-A (Ser7) antibody), methylation based on HiFi reads, and satellite enrichment were used to identify centromeric regions. The three types of repeats (LINEs in red, SINEs in brown, and satellites in light purple) are shown in the “Repeats” bar, and two satellite units (SatI and SatII) occupy the centromeric region in the “Satellites” bar. The sequence identity heatmap (bottom) with the color scale at the left bottom corner is shown across the centromere in nonoverlapping 5-kb windows, and four evolutionary layers corresponding to two layers (1 and 2) of SatI and two layers (3 and 4) of SatII in the centromeric region are marked from 1 to 4. **c,** Lengths of chromosomes (bottom) and their centromeric repeat units (top). **d,** FISH images for probes of SatIII and CenY. Two SatIII variants were identified in the T2T genome assemblies of both sheep (SatIII-20GG and SatIII-20CC) and goat (SatIII-V2A and SatIII-V2C in *T2T-goat1.0*), and their sequences were aligned below. The probes of SatIII-20GG (red) and SatIII-20CC (green) were also used for FISH imaging (right plot), while the probes of CenY (red) and SatIII (combining SatIII-20GG and SatIII-20CC, green) were hybridized onto the sheep chromosomes in FISH (left plot). The Y chromosome and CenY probes (red) in the middle of the left plot are enlarged in a white line box at the top right corner. **e**, Collinearity of three metacentric chromosomes (chromosomes 1, 2, and 3) among *T2T-sheep1.0*, *T2T-goat1.0*, and argali (*O. ammon polii*). SatI, SatII and SatIII are colored for their presence or absence in the centromeres. **f,** Phylogenetic tree of 16 species based on their SatI sequences (Supplementary Table 10).

Satellite DNAs that consisted of higher-order repeats (HORs) dominated the centromeric regions of autosomes and the X chromosome. The satellite repeats were classified into three categories: SatI (816 bp), SatII (702 bp) and SatIII (22 bp) (Supplementary Methods and Fig. 2c). SatI and SatII, which corresponded to two previously described satellites (KM272303.1 and U24091, respectively)^28,29^, dominated the centromeric regions of *T2T-sheep1.0* (Fig. 2c). We confirmed the sequence of SatI, determined the size of SatII to be 702 bp instead of the ∼400 bp reported previously^29^, and revealed a new satellite, SatIII. The centromeric SatIII repeat arrays were validated through fluorescence in situ hybridization (FISH) assays (Fig. 2d). The intensities of the FISH signals on the ends of the chromosomes were in accordance with the centromeric presence of SatIII in our assembly. Additionally, lower entropy values were found in centromeric satellites, such as, SatI and SatII on Chr02 (Fig. 2b), indicating less sequence complexity than in other chromosomal regions. The primate centromere evolved via the amplification of satellite sequences within its inner core, which forms layers of varying ages^30^. Like the presentation of centromeric regions in humans^22^, we also observed the evolutionary layers of centromeric satellites based on sequence identity and similarity heatmaps. For example, on the X chromosome, SatII dominated the centromeric region, and at least two layers were identified in the SatII HORs (Supplementary Fig. 13). Additionally, we detected the insertion of other repeat units such as LINEs and SINEs into the centromeric regions (Fig. 2b). In contrast to genes within centromeric regions, genes in pericentromeric regions were highly expressed according to the RNA-seq data (Fig. 2b).

### Evolution of centromeric satellites and chromosomal fusion

Sheep have experienced significant evolutionary events for chromosomal centric fusion of the three metacentric chromosomes of Chr1, Chr2, and Chr3^29^. Nonallelic homologous recombination (NAHR) occurred on the two acrocentric chromosomes of their wild ancestors and related species, and generated a metacentric chromosome. As a result, footprints of centromeric evolution might remain in the centromeric regions of the three metacentric chromosomes. We examined the genome of argali^5^ to trace its chromosomal reorganization using the goat genome^31^ as an outgroup (Fig. 2e). The collinearity among sheep, argali, and goat apparently revealed the two-to-one fusion relationships (CHI1 + CHI3 in goat vs. Chr01 in sheep; CHI2 + CHI8 in goat vs. Chr02 in sheep; CHI5 + CHI11 in goat vs. Chr03 in sheep) between 6 chromosomes in goat and 3 chromosomes in the two ovine species. Based on the centromeric locations on chromosomes in goats and the two ovine species, we established the chromosomal fusion pattern involving the footprints of centromeric satellites (Fig. 2e). In addition, SatII was in the core of centromeric regions with high sequence identity on the three metacentric chromosomes Chr01, Chr02, and Chr03 (Fig. 2b and Supplementary Fig. 13). According to the sequence identity heatmap of the centromeric regions of sheep, there are four evolutionary layers on Chr02 including layers #1 and #2 of SatI and layers #3 and #4 of SatII (Fig. 2b) based on identity values of 80–100%, which suggested multiple amplification events of SatI and SatII. We did not detect telomeric sequences in the centromeric regions of these three metacentric chromosomes with NAHR or fusion events in *T2T-sheep1.0*, which is different from previous observations of telomeres in centromeric regions in muntjac^32^.

We determined the sequence similarity and conservation of SatI, SatII and SatIII in Caprinae and Bovidae species, by comparing related sequences in the NCBI database. Multiple sequence alignment of SatI sequences revealed 599 variant sites among the 16 Bovidae species, and 86 variants between *T2T-sheep1.0* and *T2T-goat1.0*^31^ (Supplementary Table 10 and Supplementary Fig. 15a). The SatI sequences accumulated variants among Caprinae species, and a phylogenetic tree based on their consensus sequences showed the split of Caprinae and Hippotraginae species in the Bovidae family (Fig. 2f). The sequences of SatII harbored 130 variable sites between sheep and goat (Supplementary Fig. 15b), while two SatIII variants (SatIII-20GG and SatIII-20CC for *T2T-sheep1.0*, SatIII-V2A and SatIII-V2C for *T2T-goat1.0*) were found for both the species (Fig. 2d). As shown by the binding results of the FISH probes, the two primary SatIII variants (SatIII-20GG and SatIII-20CC) exhibited intensified signals on the chromosomal ends (Fig. 2d).

### Y chromosome structure

It has been difficult to assemble the Y chromosome due to the homology of the X and Y chromosomes^10^, i.e., pseudoautosomal regions (PARs). Thus, a relatively complete Y chromosomal assembly is available for only a few species such as human, monkey, rat, mouse, cattle, and donkey (Supplementary Fig. 16a). However, only the human^33^ (assembly *T2T-CHM13v2.0*) and six apes^15^ have a gap-free T2T Y chromosome. Hereby, *T2T-sheep1.0-chrY* of sheep was assembled and significantly improved compared with the most updated Y chromosome reference^9^ (*Ramb_v3.0-chrY*, CP128831.1) for sheep in the *Ramb_v3.0* assembly. *T2T-sheep1.0-chrY* had a length of 26.59 Mb, which was 0.67 Mb and 15.97 Mb longer than those of *Ramb_v3.0* and the earlier Hu sheep reference genome ASM1117029v1^10^ (GCA_011170295.1), respectively. The ∼17-Mb region covering the PAR showed good collinearity between *T2T-sheep1.0-chrY* and *Ramb_v3.0-chrY*, except for a 1.18-Mb inversion at ∼10 Mb (Fig. 3a). The remaining ∼9-Mb region distal to the PAR of the Y chromosome (namely Z zone) showed a low-quality assembly and highly fragmented DNA as small inversions in *Ramb_v3.0-chrY*, which was annotated as a *ZFY* gene array in *T2T-sheep-chrY* (Fig. 3a and Fig. 3b). The pairwise sequence identity heatmap showed an apparent block in this region, suggesting the presence of repeats (Fig. 3b).

**Fig. 3.**
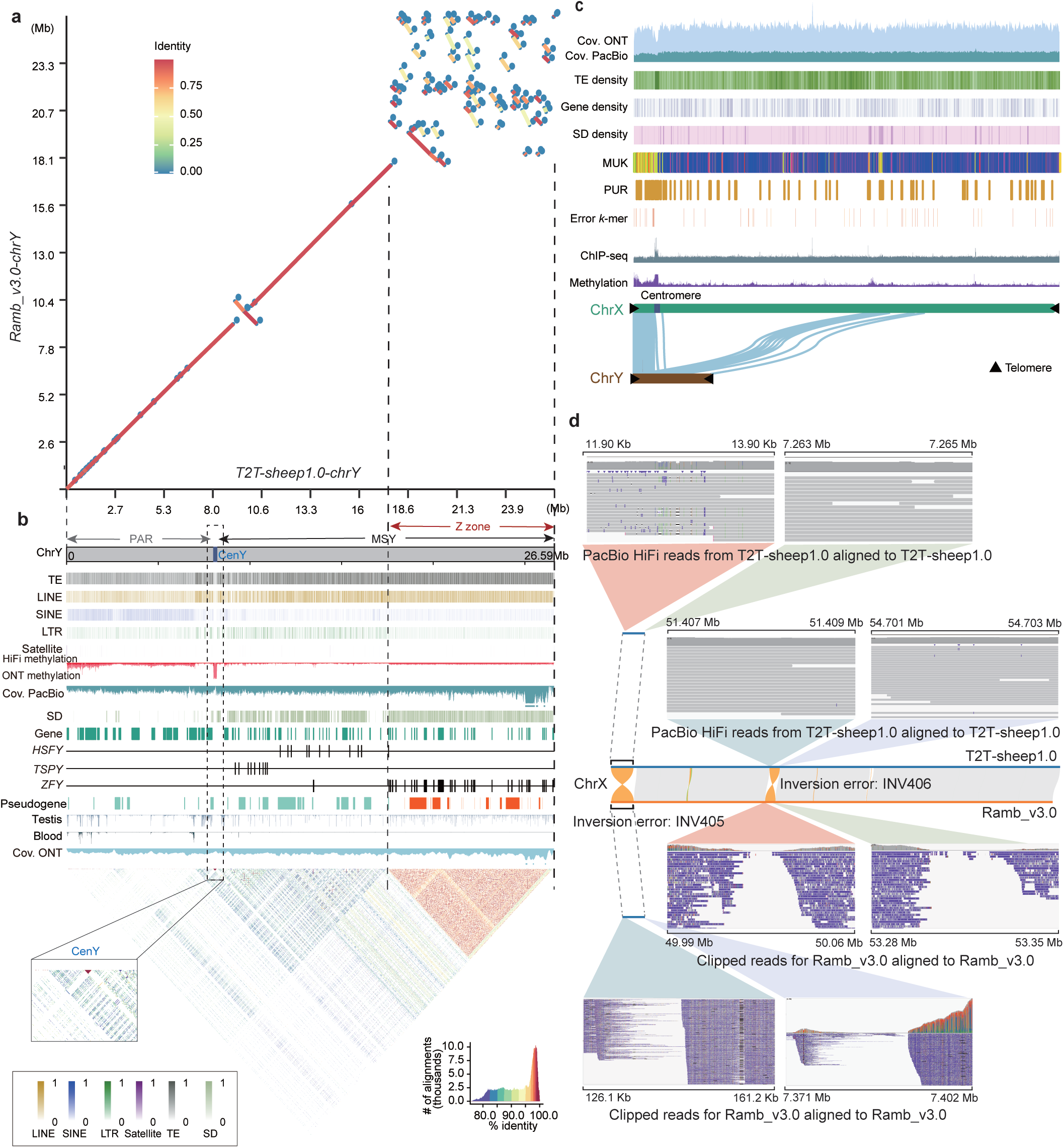
Assemblies of chromosomes X and Y. **a,** Collinearity of chromosome Y (ChrY) between *T2T-sheep1.0* and *Ramb_v3.0*, i.e., *T2T-sheep1.0-chrY* vs. *Ramb_v3.0-chrY*. The sequence identity scale is colored and placed in the top left corner. **b,** Genomic features of *T2T-sheep1.0-chrY*. From top to bottom: ChrY consists of the X-homologous region (PAR) and male-specific region (MSY), with CenY (blue) and Z zone (red) labeled; densities of TEs, LINEs, SINEs, LTRs and satellites, with scales at the bottom left corner; methylated cytosines in 10-kb windows based on HiFi (red) and ONT (light red) data; coverage of PacBio HiFi reads (Cov. PacBio); segmental duplication (SD) density in 10-kb windows; gene distribution; three multicopy genes, *HSFY*, *TSPY* and *ZFY*; pseudogenes, with the ones in the Z zone highlighted in red; Gene expression from RNA-seq of testis and blood tissues; coverage of ONT reads (Cov. ONT). Sequence identity across the whole ChrY showing the high-identity signals of CenY (magnified in the bottom left corner) and the Z zone. **c,** Genomic features annotated on chromosome X (ChrX) and regions homologous to ChrY. Homologous genes between ChrX and ChrY are connected by light blue lines. The color keys are the same as those in Fig. 1a. **d,** Two inversion errors (INV405 and INV406) on ChrX of *Ramb_v3.0* that were corrected in *T2T-sheep1.0*. The gray blocks between the horizontal bars of *T2T-sheep1.0* and *Ramb_v3.0* represent collinear regions. Orange lines indicate inversions on ChrX between *Ramb_v3.0* and *T2T-sheep1.0*. There was no coverage of Rambouillet PacBio reads that are downloaded from NCBI (Biosample ID SAMN17575729) at the two junction sites of each inversion in *Ramb_v3.0*, in contrast to the accurate assembly and uniform coverage of HiFi reads from Hu sheep in *T2T-sheep1.0*.

Centromere-specific satellites (SatI, SatII, and SatIII) were not observed, but another type of simple repeat sequence, CenY, which was 2516 bp long and spanned a total of 180.12 kb, was present on the sheep Y chromosome. As a potential centromeric repeat unit on the Y chromosome, CenY was supported by hypermethylation data and the sequence identity heatmap (Fig. 3b). Similar sequences of CenY could be detected on the goat Y chromosome^31^, but the length was only approximately 1400 bp. In addition to the Caprinae subfamily, CenY sequences are also found on the Y chromosomes of other Bovidae species, such as *Bos taurus* (GenBank accessions CP128563.1 and LR962769.1).

ChIP-seq based on phospho-CENP-A (Ser7) antibody failed to capture any reads on the whole Y chromosome of sheep, despite the presence of ChIP-seq peaks in the other chromosomal centromeric regions with SatI∼SatIII repeats (Fig. 2b and Supplementary Fig. 13). This observation is congruent with the lack of hybridization of the SatI and SatII probes to the Y chromosome observed in FISH assays of sheep^29,34^. In this study, the binding of FISH probes confirmed the unique presence of CenY on the Y chromosome (Fig. 2d). Apart from CenY, other repeats, such as LINEs, SINEs and LTRs, were also detected on the Y chromosome, accounting for 39.72%, 6.20% and 7.24% of the whole Y chromosome, respectively.

A total of 133 protein-coding genes and 59 pseudogenes were annotated (Fig. 3b). Unlike in human^33^ and goat^31^, we did not find apparent double tandem gene copies on the sheep Y chromosome, but detected significantly increased copy numbers of three gene families, i.e., 9, 11, and 33 copies for *TSPY*, *HSFY*, and *ZFY*, respectively. Our pseudogene prediction further revealed 10 more *ZFY*-like pseudogenes in the Z zone. In comparison, the *ZFY* gene on the Y chromosome showed only one copy in human *T2T-CHM13v2.0*, and 5 dispersedly distributed copies on the Y chromosome of the goat T2T genome assembly *T2T-goat1.0*^31^. Phylogenetic analysis of the *ZFY* genes confirmed their very high sequence similarity and placed them into a single clade of the tree based on nucleotide sequences (Supplementary Fig. 16b). Notably, the expansion of these three gene families was strongly related to the enrichment of SDs in these regions (Fig. 3b). RNA-seq of 148 samples covering 28 tissues from the public NCBI database confirmed the transcription of these protein-coding genes, particularly with high expression in the testis (Fig. 3b and Supplementary Fig. 17).

RNA-seq revealed no or low expression of 54 genes in the hypermethylated homologous regions in blood, compared to the abundant expression of genes in the male-specific Y (MSY) region and Z zone in the testis (Fig. 3b). *T2T-sheep1.0-chY* is one of the first complete sheep Y chromosomes with detailed gene annotation^9^, and 7 genes (i.e., *IL9R*, *IL3RA*, *SLC25A6*, *ASMTL*, *P2RY8*, *ASMT*, and *DHRSX*) on the X-chromosome-homologous region of the Y chromosome (∼4 Mb) showed an order similar to that in *human T2T-CHM13v2.0-chrY*.

### Features of the X chromosome

In addition to the Y chromosome, *T2T-sheep1.0* also significantly improved the assembly of the X chromosome, with an increase in QV from 44.76 in *Ramb_v3.0* to 51.04. The assembly of the X chromosome showed uniform coverage for ONT and HiFi reads (Fig. 3c). We corrected the mistakenly assembled inversions on the X chromosome of *Ramb_v3.0*. For example, just like INV195 on Chr09 (Fig. 1d), we confirmed errors for the two inversions (7.25 Mb for INV405 and 3.29 Mb for INV406) on the X chromosome in *Ramb_v3.0*, due to the alignment failure of PacBio reads from both Rambouillet and Hu sheep for *T2T-sheep1.0* at the two junction sites (Fig. 3d and Supplementary Fig. 9c). We annotated 959 genes on the X chromosome of *T2T-sheep1.0*, and identified centromeric regions based on enrichment of ChIP-seq and hypermethylation signals. We found that the p arm (∼7 Mb) of the X chromosome, covering 31 genes, was homologous to the ∼8.6 Mb region with 54 genes on the p arm of the Y chromosome, and was considered as PAR (Fig. 3c). Furthermore, the PAR is enriched with MUK and PURs and hypermethylated on both the X and Y chromosomes in blood (Fig. 3b and 3c). In addition, 10 genes in the middle region of the X chromosome from 81.71 Mb to 100.68 Mb exhibited collinearity with the 10 corresponding genes in the MSY region of the Y chromosome (Fig. 3c).

### Structural variants based on long reads

To investigate the performance of *T2T-sheep1.0* as a reference for calling large SVs, we sequenced the genomes of two sheep individuals from the Tan and European mouflon (Supplementary Fig. 18), and aligned their PacBio long reads to *T2T-sheep1.0* together with the downloaded datasets of the other 16 sheep samples (Supplementary Table 1 and Supplementary Table 11). The mismatch rate for alignment observed for *T2T-sheep1.0* decreased significantly (Fig. 4a), most likely because of the greater accuracy of the consensus sequences. After merging and filtering, we identified a total of 192,265 SVs overlapping with 11,987 genes (55.93% of the total genes), including 75,962 deletions (DELs) and 113,541 insertions (INSs) (Supplementary Table 12). Alignment to *Ramb_v3.0* yielded substantially less SVs across the 18 sheep samples (Fig. 4b and Supplementary Table 13). However, we discovered 663 homologous DELs and INSs with allele frequency of 36 in all 18 samples (Supplementary Table 13), less than the 959 ones with *Ramb_v3.0* as a reference, and it could be explained by structural errors corrected in *T2T-sheep1.0*. We observed a significant increase in the number of SVs in the two wild sheep of Asiatic mouflon and argali compared to the other 15 domestic sheep and European mouflon (Fig. 4b). *T2T-sheep1.0* enabled the discovery of additional 16,885 SVs within PURs spanning 24.20 Mb (Supplementary Fig. 19), most of which are deletions (*n* = 10,979) and insertions (*n* = 5473). Compared with *Ramb_v3.0*, *T2T-sheep1.0* used as the reference resulted in more deletions and insertions in highly repetitive regions with smaller size, such as satellites and SINEs, than in LINEs, LTRs, and genes (Fig. 4c and Supplementary Fig. 18). This observation can be explained by the fact that satellites dominated the PURs. In total, we observed 16,885 new SVs spanning 24.20 Mb (Fig. 4d), most of which were deletions (*n* = 10,979) and insertions (*n* = 5473). We discovered 16 SVs related to exons and homologous in all 18 individuals and their overlapping genes are related to the functions of fertility, wool, and development with a unique role in Hu sheep (Supplementary Table 14). Within the collinearity regions between *T2T-sheep1.0* and *Ramb_v3.0*, we also observed improvements in SV calling. For example, a deletion in an exon of the *TUBE1* gene was detected on Chr08 in all 18 individuals when using *T2T-sheep1.0* as a reference, and the gene assembly and annotation are supported by the presence of complete transcripts in the Iso-seq data (Supplementary Fig. 20).

**Fig. 4.**
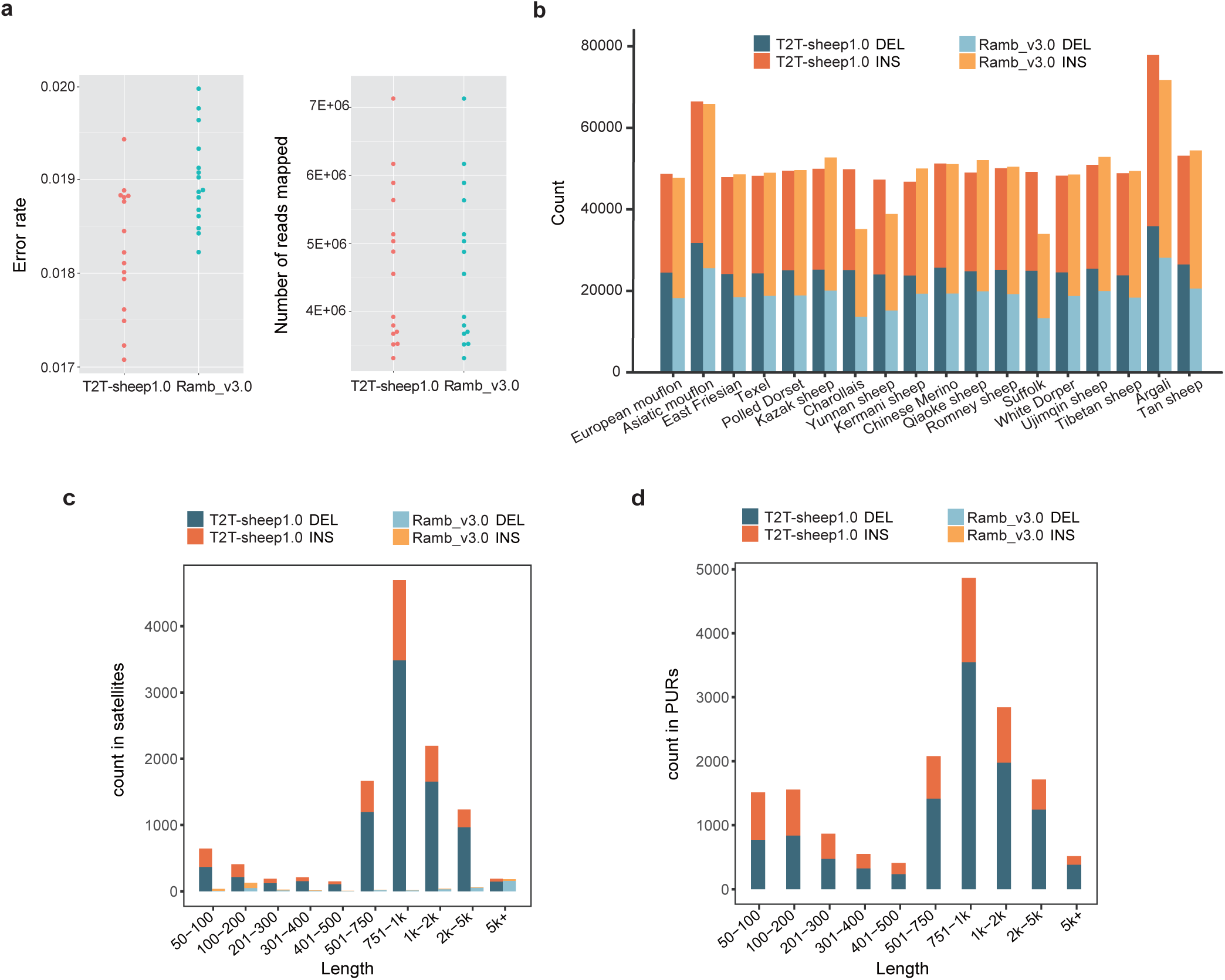
Improvements for long-read mapping and structural variant calling. **a,** Numbers of mapped reads and error rate for a ratio between mismatches and bases mapped were calculated by using Samtools based on the alignments of PacBio long reads from 18 sheep to *T2T-sheep1.0* and *Ramb_v3.0*. **b,** Deletions (DEL) and insertions (INS) derived from PacBio long reads of 18 domestic and wild sheep are compared between *T2T-sheep1.0* and *Ramb_v3.0* used as references. The counts of INSs and DELs in satellites (**c**) of *T2T-sheep1.0* and *Ramb_v3.0* and PURs (**d**) of *T2T-sheep1.0* are summed over the lengths. DEL is colored in blue for *T2T-sheep1.0* and light blue for *Ramb_v3.0*, and INS is colored in red for *T2T-sheep1.0* and orange for *Ramb_v3.0*.

### Novel genetic variants based on short reads

*T2T-sheep1.0* showed improvements in short read-based variant calling. We collected next-generation genome sequencing datasets for 810 sheep worldwide (Fig. 5a and Supplementary Table 15) and compared the SNPs detected when using *T2T-sheep1.0* and *Ramb_v1.0* as references (Supplementary Table 16). For comparison with previous results^4^, *Ramb_v1.0* was used as a reference for the alignment and call variants, rather than *Ramb_v3.0*. The depth of short reads with alignment against *T2T-sheep1.0* ranges from 10.47 to 43.41. A total of 764 (94.32%) samples showed a ≥ 10% increase in the number of mapped reads when using *T2T-sheep1.0* as a reference compared with *Ramb_v1.0*. We divided all 738 domestic sheep samples into 6 geographic populations (i.e., Central-and-East Asia, South-and-Southeast Asia, the Middle East, Africa, Europe, and America) according to their sampling locations. The remaining 72 samples comprise 7 wild species of European mouflon, Asiatic mouflon, urial, Argali, snow sheep, thinhorn sheep, and bighorn sheep. Compared to the number called by *Ramb_v1.0*, more reads were mapped to *T2T-sheep1.0* in all the geographic populations and wild species, with up to >10% added in some populations or species. We observed a much lower per-read mismatch rate when *T2T-sheep1.0* was used as the reference, while the mismatch rates of the wild species were obviously greater than those of the domestic sheep (Supplementary Fig. 21). Moreover, more reads with zero mapping quality were generated when using *T2T-sheep1.0* as the reference. This could be due to the increase in satellite sequences and SDs, which resulted in multiple locations for the alignment of short reads. Significantly fewer outward-oriented pairs were aligned with *T2T-sheep1.0*. Moreover, we detected ≥ 3% additional properly paired reads in 252 samples with alignment to *T2T-sheep1.0* (Supplementary Fig. 21). Therefore, improvements in the mapping of *T2T-sheep1.0* indicate its advantage as a sheep reference genome.

**Fig. 5.**
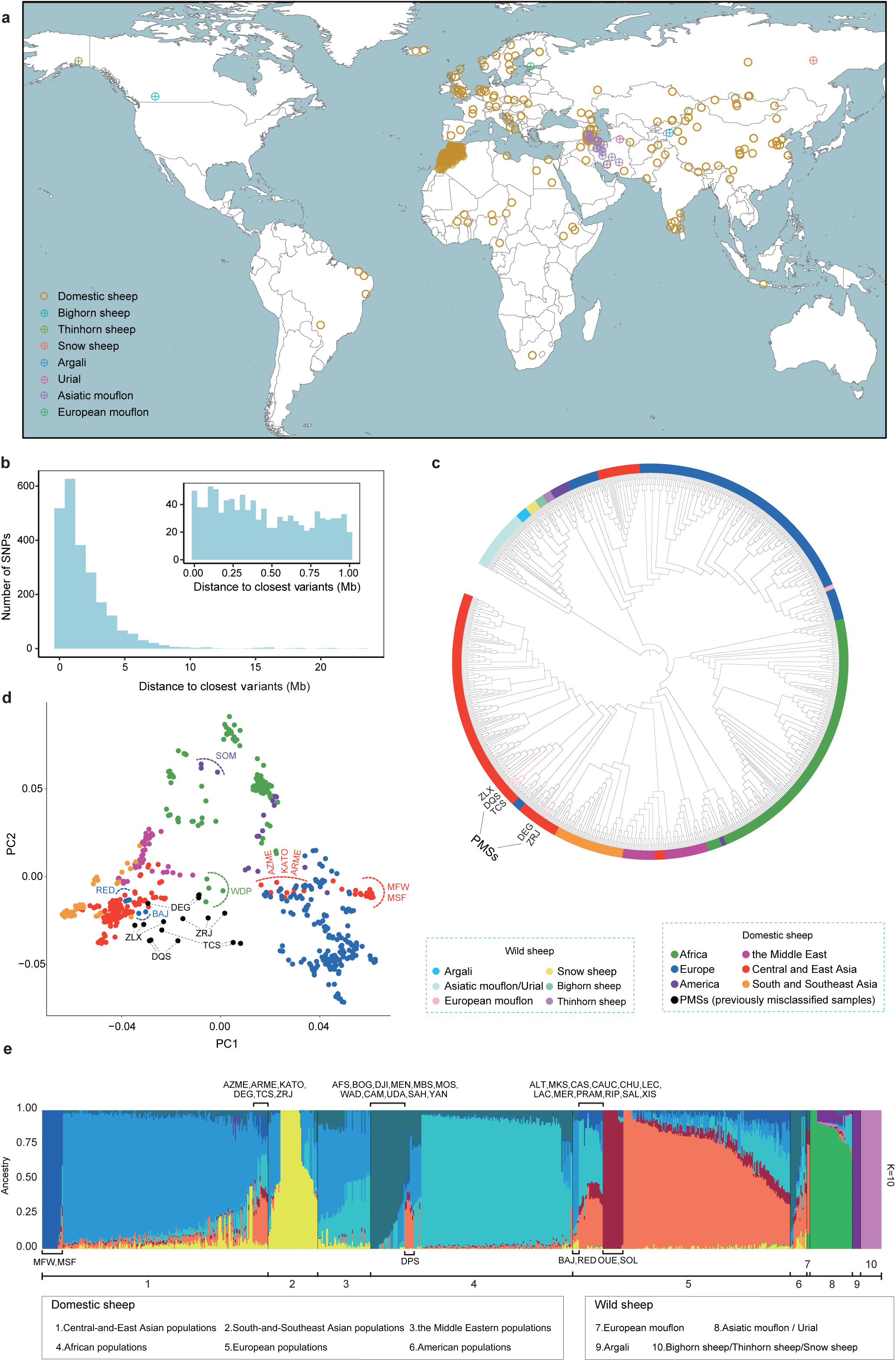
Improvement of *T2T-sheep1.0* in the analysis of short reads in sheep populations. **a,** Sampling locations of 810 NGS samples from 158 domestic sheep populations and seven wild sheep species. **b,** Distance of SNPs from PURs to the closest QTL from the AnimalQTL database. The 1-Mb distance scale is shown in the top right corner. **c,** Neighbor-joining (NJ) tree of wild and domestic sheep based on SNPs when using *T2T-sheep1.0* as a reference. Asiatic mouflon and urial genetically are not completely separated in the genetic clustering analysis and phylogenetic tree due to the presence of hybrids and gene flow between these two populations^4,60,130^. The previously misclassified samples (PMSs) are highlighted in red branches and black labels for five populations (ZLX, DQS, TCS, DEG, and ZRJ). **d,** PCA of domestic sheep populations based on SNPs using *T2T-sheep1.0* as a reference. The PMSs are highlighted in black, while the six geographic domestic sheep populations are highlighted in the same colors in both PCA and NJ tree. The populations (SOM, RED, WDP, etc.) are labeled with the colored abbreviated names because they are not clustered in the corresponding continents. **e,** Population genetic structure of wild and domestic sheep inferred from ADMIXTURE (K = 10) using *T2T-sheep1.0* as the reference. The abnormal populations in the 6 domestic sheep superpopulations (Central-and-East Asia, South-and-Southeast Asia, Middle East, Africa, Europe, and America) according to the continents are labeled with abbreviated names (Supplementary Table 15).

We obtained 133,314,255 high-quality SNP variants against *T2T-sheep1.0*, 2,664,979 of which were located in PURs (Supplementary Table 16), 12,060,995 more than observed against *Ramb_v1.0* (Supplementary Fig. 22). After further filtering SNPs with minor allele frequencies (MAFs) <0.05, 27,493,776 SNPs were used for subsequent analysis, among which 336,166 SNPs located in PURs (Supplementary Fig. 22) covering 1635 genes. Notably, we found that more low-quality SNPs were filtered with *Ramb_v1.0* as the reference, probably because of the relatively low base-level QV. The number of SNPs in the PURs on each chromosome showed no association with the length of the PURs (Supplementary Fig. 22). With *T2T-sheep1.0* as the reference, the increase in the number of total SNPs was observed in the six geographic populations and in the wild sheep population, in terms of both heterozygous and homozygous SNPs (Supplementary Fig. 22). Additionally, we identified 1,265,266 SVs (Supplementary Table 16), including 196,471 SVs in PURs, which were dominated by 1,048,080 DELs and were much more abundant than those identified in our previous study^4^ using *Ram_v1.0* as the reference.

The assembly of the PURs in *T2T-sheep1.0* provided new variants for quantitative trait locus (QTL) mapping analysis. A total of 4729 sheep QTLs related to morphological and agronomic traits were identified in 248 previous studies, according to the Animal Quantitative Trait Loci (QTL) Database (Animal QTLdb)^35^. We converted their genomic coordinates relative to *T2T-sheep1.0*, and found that 758 SNPs in the PURs were located within 2 Mb of the closest regions of the QTLs (Fig. 5b).

### Nucleotide diversity and genetic structure

SNPs called by *T2T-sheep1.0* were used to perform population analysis of wild and domestic sheep. We found the highest average nucleotide diversity *(π*) value in domestic sheep, compared to all the wild populations (Supplementary Fig. 23), while two wild sheep, Urial and Asiatic mouflon, harbored higher *π* values than previously reported for domestic sheep^4^. Phylogenetic position of sheep population is sensitive to the reference, and the analysis with *T2T-sheep1.0* as the reference resolved some samples with confusing phylogenetic positions (Fig. 5c and 5d) in the neighbor joining (NJ) tree and principal component analysis (PCA). In the NJ tree with *Ramb_v1.0* as the reference, five populations originating from Southwest China (Diqing with a label of DQS, Tengchong with TCS, and Tibetan sheep with ZRJ and ZLX) and Kazakhstan (Degeres mutton-wool sheep with DEG) were not placed in the clade of Central and East Asia (Supplementary Table 15), while the NJ tree with *T2T-sheep1.0* was updated with these five populations in the Central and East Asian clade. So we labeled them as previously misclassified samples (PMSs), and *F*_ST_-based Neighbor-Net network further confirmed close phylogenetic relationships of PMSs with Central and East Asian sheep (Supplementary Fig. 24).

Genetic structure by ADMIXTURE (*k*=10) and *F*_ST_-based Neighbor-Net network based on the SNPs showed consistent patterns of genetic differentiation among domestic (six populations: red for Europe, green-blue for Africa, light-blue for Central-and-East Asia, yellow for South-and-Southeast Asia, and mosaic colors for the Middle East and America) and wild (four populations) sheep populations according to their geographic origins (Fig. 5c, 5d, 5e, and Supplementary Fig. 24). Furthermore, genetic divergence of the lineages was observed inside the domestic sheep populations on the continents. For example, Chinese Merino (abbreviated as MFW and MSF) and six breeds of Central Asia and Tibet (AZME, ARME, KATO, DEG, TCS, and ZRJ) in Central-and-East Asia received the genetic introgression of European sheep with closer relationships with European clade (Supplementary Fig. 24) and showed the colors of blue and red respectively, rather than light-blue (Fig. 5e). African sheep consist of two groups (green-blue and dark green-blue in Fig. 5e) and contain a breed of Dorper sheep (DPS in orange in Fig. 5e) with European blood. European sheep have the genetic introgression of African sheep in 12 breeds of South Europe (ALT, MKS, etc.) in light-blue, while North European sheep (OUE and SOL) also show the different lineage origin.

### Selection signatures for domestication

To confirm the improved ability of *T2T-sheep1.0* to identify genomic regions selected during domestication, we reanalyzed the sequencing data in a genomic comparison between Asiatic mouflon and five old domestic landrace populations from a previous study in which the sheep assembly *Oar_v4.0* (GCA_000298735.2) was used as a reference^36^. The regions with the top 1% of outliers for the cross-population composite likelihood ratio (XP-CLR) and *F*_ST_ were considered as candidate selective sweeps. A total of 1066 selected regions of 53.30 Mb and covering 338,024 SNPs and 197 SVs were identified with extreme allele frequency differentiation across the 27 chromosomes (Supplementary Table 17 and Supplementary Table 18). Overall, 311,888 SNPs (92.27%) associated with the top 1% selected regions as detected with *T2T-sheep1.0* could be successfully mapped onto *Oar_v4.0*, and 1403 genes within these sweeps were designated candidate selected genes (Fig. 6a). We discovered multiple novel selection signals (blue-colored in Fig. 6a) in the PURs of pericentromeric regions, such as those on Chr03, Chr17, Chr18, and Chr24. A total of 146 candidate genes obtained using *Oar_v4.0* as a reference^36^, such as *MBOAT2*, *TEX12*, *PDE6B* and *CUX1*, were also included in the list of selected genes detected by *T2T-sheep1.0* (Fig. 6b).

**Fig. 6.**
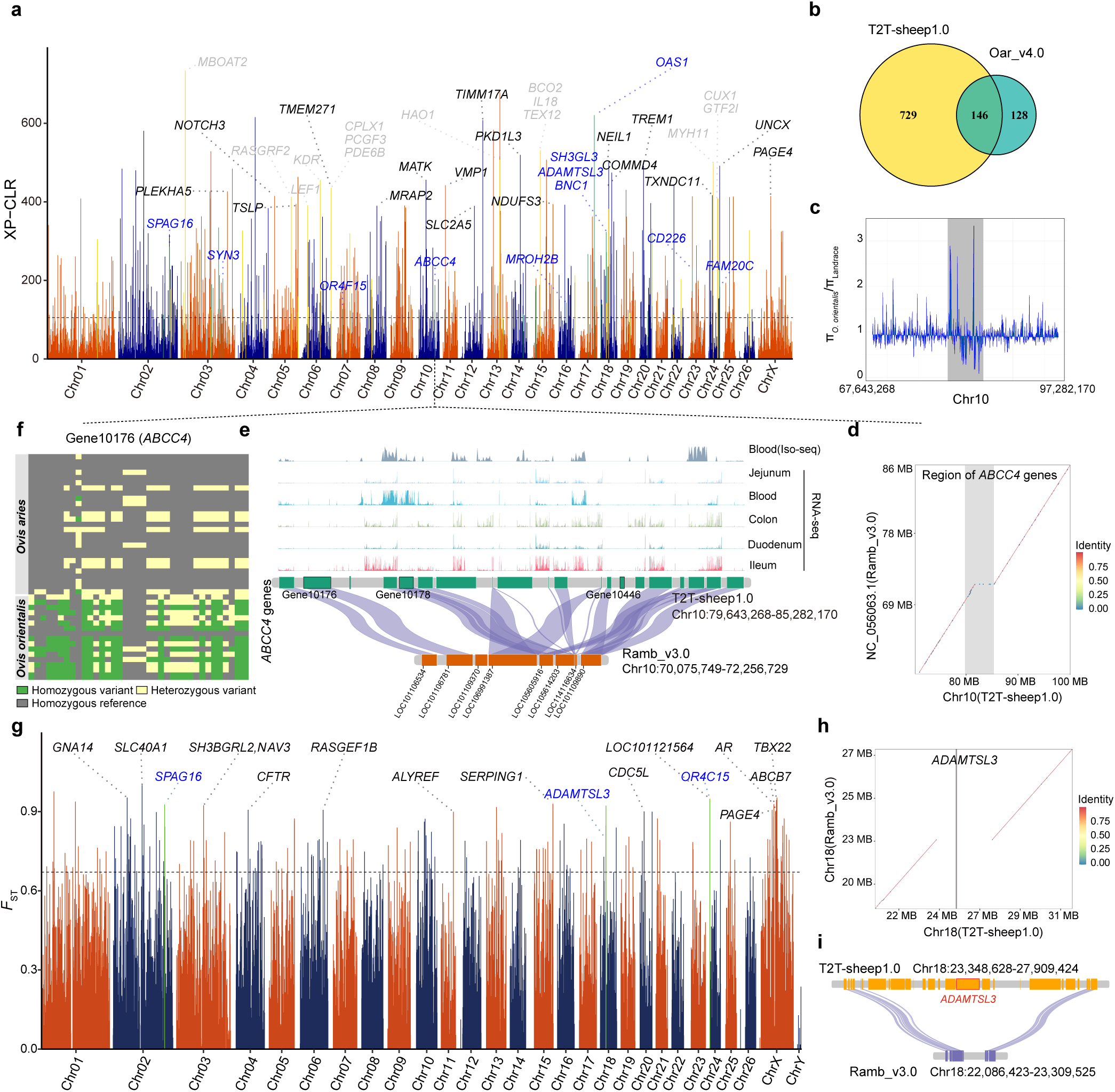
Selection signatures associated with domestication. **a,** Selection signals based on SNPs and the top 1% of XP-CLR values (horizontal dash line) for landrace sheep breeds compared with wild sheep of Asiatic mouflon using *T2T-sheep1.0* as a reference. The genes identified in non-PURs by both *T2T-sheep1.0* and *Oar_v4.0* are shown in gray, the ones in non-PURs identified only by *T2T-sheep1.0* in black, and the ones in PURs in blue. **b,** Venn diagram of selected genes associated with domestication based on SNPs between *T2T-sheep1.0* and *Oar_v4.0* as references. **c,** The π ratio (π-O. orientalis/π-landrace) confirms strong selection signals detected by XP-CLR values in the region of the *ABCC4* gene family on Chr10. **d,** The collinearity between the two assemblies of *T2T-sheep1.0* and *Ramb_v3.0* showing a PUR that corresponded to the selected region with the *ABCC4* gene family is highlighted with a gray bar. **e,** Twenty *ABCC4* family genes are included in the selected region on Chr10, while only eight ones were assembled in *Ramb_v3.0*. The *ABCC4* genes are transcribed in blood, jejunum, colon, duodenum, and ileum tissues, as shown in RNA-seq analysis. The three *ABCC4* genes under domestication selection are highlighted with black outlines. **f,** Thirty-seven nonsynonymous variants of the *ABCC4* gene “Gene10176” with differences between domestic and wild sheep. **g,** Selection signals based on SVs and top 1% of *F*_ST_ values assessed between landrace and wild sheep with *T2T-sheep1.0* as a reference. **h,** A selected SV on Chr18 was located in PURs, and the collinearity between *T2T-sheep1.0* and *Ramb_v3.*0 confirmed the presence of this PUR containing the *ADAMTSL3* gene. **i,** The ∼3.5-Mb region with the *ADAMTSL3* gene has been wrongly assembled from Chr18 of *Ramb_v3.0* and considered a PUR in *T2T-sheep1.0*.

In particular, 550 and 36 novel selected genes that were not identified by *Oar_v4.0* were identified in non-PURs (e.g., *TIMM17A, PLEKHA5, UNCX* and *PKD1L3*; gray-colored in Fig. 6a) and PURs (e.g., *ABCC4*, *SPAG16*, *OAS1*, *BNC1*, *CD226* and *FAM20C*; blue-colored in Fig. 6a) of *T2T-sheep1.0*, respectively (Supplementary Fig. 25 and Supplementary Table 17). These novel genes were mostly involved in immunity, neuron development, sperm, energy metabolism, etc. For example, we detected selective signals of a ∼4 Mb region (Chr10: 80,150,000-83,900,000) by both XP-CLR and nucleotide diversity (π) ratio of π-O. orientalis/π-landrace (Fig. 6c). This selected region covered 20 *ABCC4* gene copies (Gene10176∼10179, Gene10434, Generf10555, Gene10437∼10438, Generf10560, Gene10440, Generf10562, Gene10443, Generf10563, Gene10445∼10446, Gene10448∼10452, Fig. 6d) and was not assembled in *Ramb_v3.0* with only 8 truncated short *ABCC4* gene copies (Fig. 6e). These 20 *ABCC4* copies were expressed in multiple tissues, including blood, colon, duodenum, and ileum, indicating specialized functions. Further examination revealed 37 nonsynonymous mutations in one *ABCC4* copy (Gene10176, Fig. 6f) and selected sites in the other two *ABCC4* copies (Gene10178 and Gene10446) in domestic sheep compared to wild sheep (Fig. 6e and Supplementary Fig. 25a). The selection signals of other novel genes (e.g., *SPAG16*, *OAS1*, *BNC1*, *CD226* and *FAM20C*) were also confirmed based on the π ratio and differentiated alleles between the wild and landrace sheep populations (Supplementary Fig. 25b).

Seven SVs within the top 1% of the *F*_ST_ outliers were identified in the PURs (Fig. 6g and Supplementary Table 18). Within an ∼3.5-Mb PUR (Fig. 6h), we identified two deletions (1.06 kb and 1.49 kb in length) located within the introns of the *ADAMTSL3* gene on Chr18 (Supplementary Fig. 25c), which plays a cardioprotective role in maintaining cardiac function in human and mice^37^. This PUR in *T2T-sheep1.0* included 24 genes not present on Chr18 of *Ramb_v3.0* (Fig. 6i). A newly identified deletion in one intron of *SPAG16*, which is involved in the development, maturation, and motility of sperm^38^, was also detected in a small PUR, and allele frequencies of this deletion and other SNPs revealed significant differentiation between five landrace sheep breeds and Asiatic mouflon (Supplementary Fig. 25d).

### Selection signatures for fleece fiber diameter

We applied *T2T-sheep1.0* to detect genome-wide selection signatures among hairy and coarse-, medium-and fine-wool domestic sheep populations with decreasing fleece fiber diameters based on both SNPs and SVs (Supplementary Table 19 and Supplementary Table 20). To compare the results, we followed the same analysis procedures used in our previous study, with *Ramb_v1.0* as a reference^4^. The top 1% of XP-CLRs revealed 1014 selection signals between fine-wool and hairy sheep, and 383,248 (98.54%) SNPs within the top 1% of the selected regions based on *T2T-sheep1.0* could be successfully lifted over to *Ramb_v1.0*, and 228 genes, including *TP63*, *KRT* (*KRT77*, *KRT1*, *KRT2*, *KRT74* and *KRT71*), and *IRF2BP2*, were shared when using two reference genomes (Fig. 7a). These genes known to be under selection^4^ were confirmed based on the selected sites in domestic sheep (Supplementary Fig. 26a). Compared to those in *Ramb_v1.0*, ∼779 and 24 novel selected genes were identified in non-PURs and PURs, respectively, in the comparison of fine-wool and hairy sheep (Fig. 7b). For example, *TARBP1*, *EPS8*, and *DMXL2* in non-PURs were identified with selected alleles for the fine-wool trait (Supplementary Fig. 26b). *FOXQ1* was identified in a PUR at the end of Chr20, whose selection is supported by the π ratio between fine-wool and hairy sheep (Fig. 7c), and *FOXQ1* was reported to play a role in hair follicle differentiation^39^. The incomplete and misassembled end of Chr20 in *Ramb_v3.0* was confirmed in the collinearity analysis with *T2T-sheep1.0* (Fig. 7d). We explored the variants in *FOXQ1*, and found five variants with different allele frequencies between the coarse-, medium-and fine-wool sheep populations and the hairy population (Fig. 7e). Moreover, we detected significant selection signatures in *FOXQ1* in the other three comparisons of hair vs. coarse wool, hair vs. medium wool, and fine wool vs. medium wool (Fig. 7e and Supplementary Fig. 27).

**Fig. 7.**
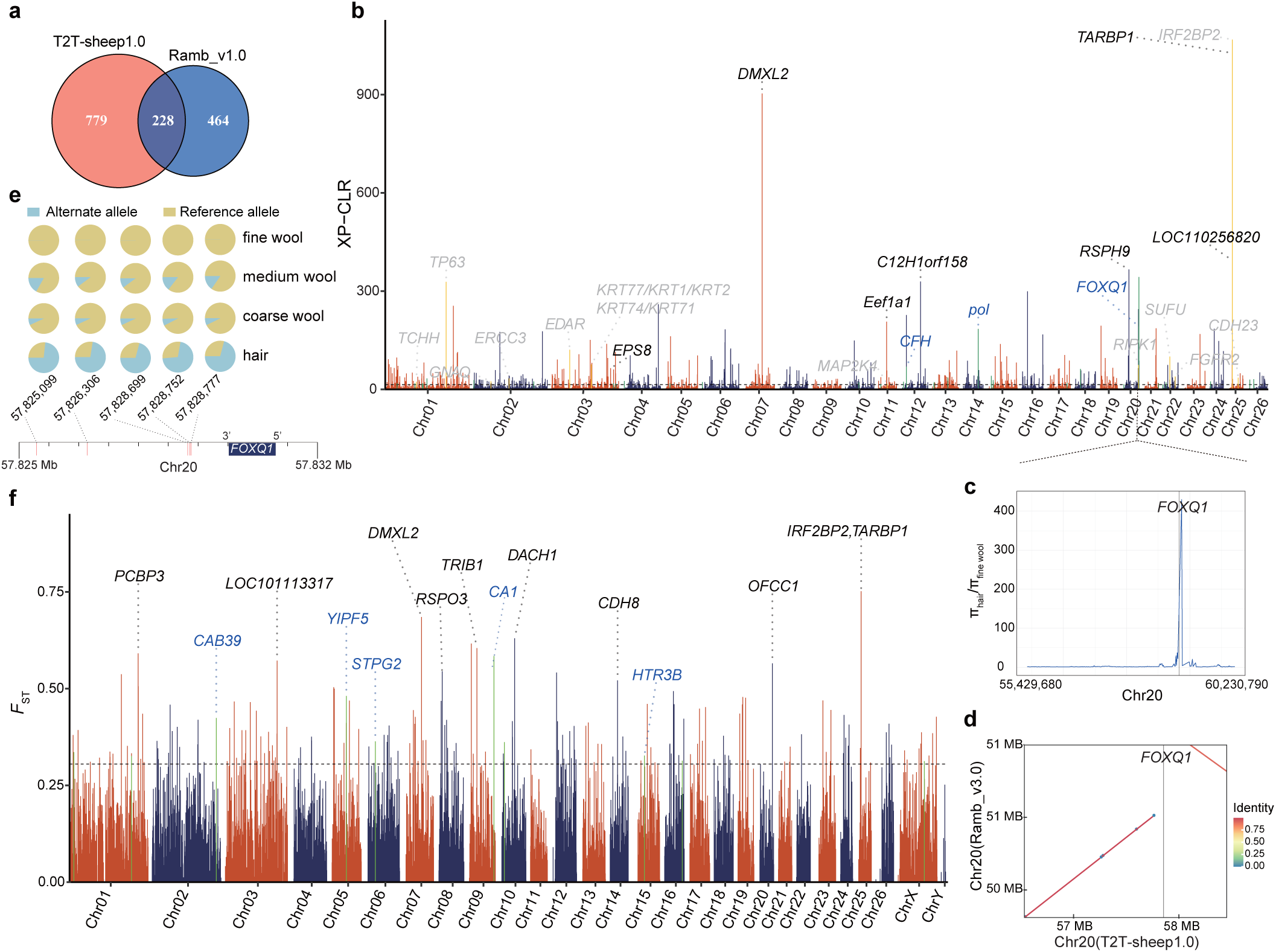
Selection signatures associated with fleece fiber diameter. **a,** Venn diagram of selected genes for fine-wool sheep based on SNPs between *T2T-sheep1.0* and *Ramb_v3.0* used as references. **b,** Selection signals based on SNPs and the top 1% of XP-CLR values when comparing fine-wool and hairy sheep. **c,** The π ratio (π-hair/π-fine wool) between hairy and fine-wool sheep confirmed selection on the *FOXQ1* gene on Chr20 detected by the XP-CLR. **d,** The selected region with the *FOXQ1* gene in gray is located in a PUR at the right end of Chr20 in *T2T-sheep1.0* compared to *Ramb_v3.0*. **e,** Five mutation sites at the 3’ downstream region of the selected *FOXQ1* gene and their allele frequencies showed selection in coarse-, medium- and fine-wool sheep, compared to the hairy sheep population. **f,** Selection signals based on SVs and the top 1% of *F*_ST_ values detected between fine-wool and hairy sheep.

We also identified 195 candidate SVs on the basis of the top 1% of *F*_ST_ values between the fine-wool and hairy populations when using *T2T-sheep1.0* as a reference (Supplementary Table 20). The strongest signal was derived from an insertion in 3’ UTR of the *IRF2BP2* gene located on Chr25, and the inserted fragment was previously identified and determined as an antisense *EIF2S2* retrogene (called as *EIF2S2*)^40^. The selection of *IRF2BP2* gene is also supported by eight SNP sites with significant allele differences between fine-wool and hairy populations, including two SNPs in the 3’ UTR and intron of *IRF2BP2* gene in our previous findings^4^, and six more SNPs in the promoter (<2000 bp away from the transcription initiation site) and ∼5 kb upstream regions in this study (Fig. 7f and Supplementary Fig. 28). Another top signal detected based on both SNPs and SVs revealed a deletion within the intron of *DMXL2* (Supplementary Fig. 26b and Supplementary Fig. 26c). Both the SNPs and SVs of these two genes have been under selection due to obvious differences between the hairy and fine-wool populations (Fig. 7b and Fig. 7f). Some genes overlapping with selected SVs, such as *RSPO3*^41^ and *OFCC1*^42,43^, reportedly have functions related to hair follicle development and hair curling. Besides, nine selected SVs overlapped with PURs. For example, a deletion (1763 bp) under selection was located in the intron of *CA1* within a PUR (Supplementary Fig. 26d). Selective signals for *CA1* were also found in three other comparisons: fine-vs. coarse-wool, medium-vs. coarse-wool, and medium-wool vs. hairy sheep (Supplementary Fig. 29). We used PacBio long reads to verify the SVs in PURs called by short reads, and five of the 7 SVs associated with domestication and all 9 SVs associated with selection for wool fineness trait were confirmed (Supplementary Table 21).

## Discussion

Since the release of human *T2T-CHM13*, T2T genome assemblies have become popular and available for several species^16,44,45^. Nevertheless, several gaps are still present in recent nearly complete animal genome assemblies, including those of chicken^46^, duck^47^, Mongolian gerbil^48^, and cattle^49^. The *T2T-sheep1.0* genome assembled here represents the first gap-free T2T genome of a ruminant, which we obtained by resolving PURs, particularly those in centromeres and on the Y chromosome (*T2T-sheep-chrY*).

In earlier years, different assembly strategies have been adopted for the Y chromosome, including the utilization of third-generation long-read sequences for sheep^10^, and BACs and Y-chromosomal markers for human^50^, threespine stickleback^51^, and horse^52^. Quite recently, Y-chromosome assemblies of human and brown planthopper^14,33,53^ were performed with the newly developed method of trio-binning haplotype-resolved assembly and parental sequencing data^12^. *T2T-sheep-chrY* was assembled with the recent approach as above, and became the representative Y assembly of the family Bovidae. Multiple copies of the *TSPY*, *HSFY*, and *ZFY* genes were detected in *T2T-sheep1.0*. Multiple copies of spermatogenesis-related genes on Y chromosome promote healthy sperm function, ensure the proceed of spermatogenesis in spite of loss of some copies, and mitigate further gene loss in male animals^54^. In human, multiple copies of the *TSPY*, *DAZ*, and *RBMY* genes were detected on the Y chromosome^33^. Also, a growing body of evidence showed that multiple copies of the *TSPY* gene have been reported on Y chromosome of other animals, such as goat (26 copies)^31^ and cattle (94 copies)^55^, which could be explained by the regulation of spermatogenic efficiency via highly variable copy dosage^33^.

The *ZFY* gene is also involved in spermatogenesis and is potentially a sex determination factor in Hu sheep^56^. Two *ZFY* genes (*ZFY1* and *ZFY2*) in mice and only one *ZFY* gene with two major splice variants in human are essential for sperm formation^57^, and only five diversely-distributed *ZFY* genes were identified on the goat Y chromosome^31^. In contrast, the bovine Y chromosome harbors 136 *TSPY*, 192 *HSFY*, and 313 *ZFY* genes (including 79 *ZNF280AY* and 234 *ZNF280BY* genes)^58^. In this study, the amplification of *ZFY* genes occurred in the Z zone distal to the PAR of *T2T-sheep-chrY* (Fig. 3a), and a similar structure of the satellite (HSat) arrays was also discovered at the distal end of the q arm on the human Y chromosome^33^. The similar phenomenon for satellite expansion distal to the PAR of T2T-level Y chromosome was also observed in the apes^15^. It is believed that the strategy of repeats on the distal end of Y chromosome might be functional and beneficial to the maintenance of Y chromosome in male animals. One possible hypothesis is of a potential role in preventing the recombination between chromosomes X and Y and avoiding loss of male-specific genes on Y chromosome during crossover of meiosis^59^. The chromosomal structure, centromeric location, ampliconic genes, and multiple copies of *HSFY*, *TSPY* and *ZFY* genes in *T2T-sheep-chrY* were confirmed by those in *Ramb_v3.0-chrY*^9^. However, *Ramb_v3.0-chrY* is 0.69 Mb shorter than *T2T-sheep1.0-chrY*, which is primarily attributed to the highly fragmented assembly of the Z zone on *Ramb_v3.0-chrY* (Fig. 3a). Compared with other sheep assemblies^7,10,60^, *T2T-sheep1.0* with the addition of the complete Y chromosome may facilitate sheep genomic studies involving rams, for example, paternal lineage analysis with improved alignments.

Centromeric sequences provide evidence for chromosomal evolution in sheep. Consistent with the pattern observed for the human assembly *T2T-CHM13*, the centromeric regions in *T2T-sheep1.0* dominated the PURs. Contrary to the traditional view of gene-poor centromeres, centromeric regions in *T2T-sheep1.0* contain many genes, albeit with no or extremely low levels of expression across multiple tissues. The binding of centromeric proteins (e.g., CENP-A, CENP-C, and CENP-E) inhibited transcriptional initiation^61^. Due to the lack of known transcripts and proteins, it is unknown whether these centromeric genes are functional, but their mutations and evolution are worth investigating in the future. Given that these genes are still intact, it is very likely that they are under selective constraint and thus still are functional. In rice, ∼41% of 395 non-TE genes that were found in centromeric regions are transcribed^62^. However, no centromeric genes have been reported in human *T2T-CHM13*^13^, but they can be found in neo-centromeres for centromere repositioning^61^ or in the pericentromeric regions^26^. Additionally, we discovered four centromere repeat units and their variants, including two new ones SatIII and CenY, and two known ones, SatI and SatII^29,34^. While many previous studies estimated sheep SatII sequences to be approximately 400 bp^63^ in length, the *T2T-sheep1.0* assembly shows that its length actually is 702 bp. Conservation of SatI sequences among the Bovidae species was confirmed by the phylogenetic tree built from *p*-distance for nucleotide differences (Fig. 2f), as described previously for the SatI sequence analysis in Bovidae^64,65^. The similarities of sheep centromeric repeats retained the footprints of centric fusions or Robertsonian translocations on Chr01, Chr02, and Chr03, as the satellite DNA sequences are believed to promote chromosomal rearrangements and NAHR^64^. CenY with a size of 2516 bp was uniquely present on *T2T-sheep-chrY*, as confirmed by FISH (Fig. 3b), a phenomenon commonly reported in other mammals. For example, a unique centromeric repeat unit (1747 bp) of the Y chromosome was discovered in gerbil^48^. Also, 34-mer HORs of alpha-satellites are observed in the Y chromosome of human *T2T-CHM13*, differing from the centromeric alpha-satellites of 171-bp monomer in autosomes^33^.

The improvement of *T2T-sheep1.0* provided more accurate chromosome sequences, by correcting structural and base-level errors in the current reference genome *Ramb_v3.0*, and resulted in more variants being detected than in previous studies^4,36^. A total of 170,396 SVs (with 149,158 deletions and insertions) were found in the pangenome of 15 individual sheep genomes^66^, while in this study, 192,265 SVs (with 189,503 deletions and insertions) were discovered with *T2T-sheep1.0* as a reference in long reads from 18 individuals. The complete assembly of repetitive sequences in *T2T-sheep1.0* enabled us to identify additional duplicated genes and variants related to domestication and selection on wool fineness, whose copies or duplications inhibited accurate assembly in the previous assemblies (Supplemental Fig. 12). For example, *ABCC4* genes within a selected region associated with domestication formed a duplicated gene cluster on Chr10, and the SDs inhibited accurate assembly and generated a gap in the previous assemblies. Compared with that in wild goats, the copy number of *ABCC4*, a gene which was involved in the immune response, decreased in domestic goats^67^. As gene duplication commonly occurs under selection in the sheep genome^68^, a selective PUR associated with fleece fiber diameter contains three FOX family genes (*FOXQ1*, *FOXF*, and *FOXC*) at the end of Chr20. We also identified some more alleles for selection of genes, for example, six more SNP sites of *IRF2BP2* gene associated with fleece fiber phenotype (Supplementary Fig. 28), in addition to the three previously identified variations^4,40^. The LD analysis showed strong linkage of these 9 alleles of *IRF2BP2*, probably being responsible as a whole for the selection of *IRF2BP2* (Supplementary Fig. 28).

Our study shows that *T2T-sheep1.0* provides a gap-free complete sheep genome, and this assembly is expected to promote more comprehensive research on genome evolution, the detection of SVs and SNPs, and the discovery of gene functions in sheep and related species.

## Methods

### Sample collection

We selected Hu sheep, a popular Chinese native breed with high fertility and unique white lambskin^69^, for the T2T genome assembly. Blood from a 4-month-old ram (HU3095) and its parents were collected at the Qianbao Animal Husbandry Co., Ltd. (latitude 33.4761° N, longitude 120.2795° E, Yancheng City, Jiangsu Province, China), using 10 ml BD Vacutainer blood collection tubes (Cat. No. 368589, Becton Dickinson, NY, USA) with EDTA. The blood was then stored in a −80°C freezer before the extraction of RNA and DNA or formaldehyde cross-linking for sequencing. Blood from one adult ewe of Tan sheep (a Chinese native sheep breed well-known for its pelt) and an adult ewe of European mouflon (*Ovis musimon)* were sampled, followed by DNA extraction and PacBio sequencing. Long-read sequences of the two sheep samples, together with those of 16 sheep downloaded from the NCBI database (Supplementary Table 1), were included in the SV calling. All experimental protocols in this study were reviewed and approved by the Institutional Animal Care and Use Committee of China Agricultural University (CAU20160628-2).

### DNA extraction, long-read sequencing, and optical map generation

For HU3095, high quality genomic DNA for sequencing was extracted by the cetyl-trimethylammonium bromide (CTAB) method and purified with a QIAGEN^®^ Genomic Kit (Cat#13343, QIAGEN, Beijing, China). The 20-kb single-molecule real-time sequencing (SMRT) bell libraries for HU3095 (i.e., the T2T assembly individual) and 15-kb libraries for the two other individuals (i.e., Tan sheep and European mouflon) were prepared using the SMRTbell Express Template Prep Kit 2.0 (Pacific Biosciences, CA, USA) and sequenced on a PacBio Sequel II system according to the standard protocol (Pacific Biosciences, CA, USA).

Ultralong Nanopore libraries were constructed with approximately 10 μg of size-selected (>200 kb) genomic DNA with the SageHLS HMW library system (Sage Science, MA, USA), and then processed using a Ligation Sequencing Kit (Cat# SQK-LSK109, Oxford Nanopore Technologies, Oxford, UK) following the manufacturer’s instructions. DNA libraries (approximately 800 ng) were sequenced on a PromethION instrument (Oxford Nanopore Technologies, Oxford, UK).

Additionally, ultra-high-molecular weight (uHMW) DNA was extracted from fresh blood using a modified Bionano Prep Blood DNA Isolation Protocol (Cat# 30033, Bionano Genomics, San Diego, CA, USA), and labeling and staining were performed with DLE-1 enzyme (Bionano Genomics) according to the Bionano Prep Direct Label and Stain (DLS) protocol (Cat# 30206, Bionano Genomics). Stained DNA was loaded onto Saphyr chips and imaged in the Bionano Genomics Saphyr System (Bionano Genomics).

### Short-read sequencing

The whole-genome sequencing libraries for short reads were prepared using an MGIEasy FS DNA Prep Kit (MGI, Shenzhen, China), and 150-bp paired-end sequencing was performed on the DNBSEQ-T7RS platform (MGI) for sample HU3095 and its parents. Hi-C libraries were prepared from cross-linked chromatin of white blood cells according to a previous Hi-C protocol^70^ and sequenced on a DNBSEQ-T7RS platform (MGI).

Chromatin immunoprecipitation sequencing (ChIP-seq) was conducted to identify the centromeric regions using phospho-CENP-A (Ser7) rabbit polyclonal antibody (Cat# 2187, Cell Signaling Technology, Beverly, MA, USA). Approximately 10 ml of fresh blood was collected from the sheep HU3095, and nucleic DNA was extracted and crosslinked in 1% formaldehyde for 15 min. The crosslinking reaction was quenched with 200 mM glycine, and the DNA[protein complex was sonicated using a Covaris E220 Focused-ultrasonicator (Woburn, MA, USA). For ChIP-seq, chromatin was incubated with the phospho-CENP-A (Ser7) antibody mentioned above for DNA purification. Libraries were constructed following the instructions of the Illumina ChIP-seq Sample Prep Kit (Cat# IP-102-1001, Illumina, San Diego, CA, USA) and sequenced to generate 150-bp paired-end reads on the Illumina NovaSeq-6000 platform.

### RNA extraction, RNA-seq and Iso-seq

We performed RNA-seq and Iso-seq of HU3095 for subsequent genome annotation. Total RNA was isolated from the blood of HU3095 (Supplementary Table 1) using an RNAprep Pure Tissue Kit (Cat# 4992236, TIANGEN Biotech, Beijing, China) according to the manufacturer’s instructions. High-quality RNA samples (RIN>8, OD260/OD280=1.8–2.2, OD260/OD230>2.0) were used to construct the RNA-seq and Iso-seq libraries. For RNA-seq, RNA was first fragmented into small pieces using fragmentation reagents from the MGIEasy RNA Library Prep Kit V3.1 (Cat# 1000005276, MGI). The first strand of cDNA was then synthesized using random primers and reverse transcriptase, followed by second-strand cDNA synthesis. Based on double-stranded cDNA, short-read RNA-seq libraries were prepared using the MGIEasy RNA Library Prep Kit V3.1 (Cat# 1000005276, MGI), and sequenced on the DNBSEQ-T7RS platform (MGI) to generate paired-end reads. For gene annotation and tissue-specific expression analysis, 148 RNA-seq datasets from 28 tissues of Hu sheep, were downloaded from the NCBI database (Supplementary Table 7).

For Iso-seq of HU3095, cDNA was synthesized using polydT primers and a SMARTer PCR cDNA Synthesis Kit (Cat# 634926, TaKaRa Bio, Shiga, Japan), and double-stranded cDNA was synthesized via downstream large-scale PCRs using PrimeSTAR GXL DNA Polymerase (Cat# R050A, TaKaRa Bio, Shiga, Japan). Full-length cDNAs were used to construct sequencing libraries with the SMRTbell Template Prep Kit 2.0 (PacBio Biosciences), and sequenced on the PacBio Sequel II platform using the Sequel Binding Kit 2.0 (PacBio Biosciences).

### Fluorescence in situ hybridization (FISH)

FISH was performed as previously described with minor modifications^5^. The 2516-bp CenY and 22-bp Sat III sequences were synthesized and labeled with Dig-dUTP or Bio-dUTP (Roche Diagnostics, Basel, Switzerland) using Nick Translation Mix (Roche, Mannheim, Germany). Chromosome preparations were made from fibroblast cultures derived from skin biopsies. Slides with cell suspensions at metaphase were hybridized with a hybridization mix containing probes, and the hybridization was detected with signals of Alexa Fluor 488 streptavidin (Thermo Fisher Scientific, Waltham, MA, USA) for biotin-labeled probes and rhodamine-conjugated anti-digoxigenin (Roche Diagnostics, Basel, Switzerland) for digoxigenin-labeled probes. Chromosomes were counterstained with DAPI (Vector Laboratories, Odessa, Florida, USA). FISH images were observed using an Olympus BX63 fluorescence microscope equipped with an Olympus DP80 CCD camera (Olympus, Tokyo, Japan).

### Long-read and short-read sequence data for SV and SNP calling

To assess the performance of *T2T-sheep1.0* as a reference genome for mapping long reads and calling SVs, we collected PacBio HiFi and PacBio CLR data from 16 sheep individuals from the NCBI database and generated long reads from one Tan sheep and one European mouflon (Supplementary Table 11). In total, we obtained long-read datasets of 3 wild sheep (Asiatic mouflon, *O*. *orientalis*: argali, *O*. *ammon*: and European mouflon, *O. musimon*) and 15 domestic sheep individuals representing 15 breeds worldwide.

Additionally, whole-genome short-read sequences of 810 samples representing 72 wild (including 32 Asiatic mouflon, 6 bighorn sheep, 6 thinhorn sheep, 9 urial, 8 argali, 8 snow sheep, and 3 European mouflon) and 738 domestic sheep from 158 populations were retrieved from public databases. The short-read sequences showed an average sequencing coverage of 16.1× and were included in the variant calling for population genomics analyses, including population structure, phylogenetic tree, and selection signature detection (Supplementary Table 15).

### Initial assembly based on HiFi reads

PacBio HiFi reads were used to construct the initial assembly of autosomes and the X chromosome after removing low-quality reads and adapters. HiFi reads were generated using circular consensus sequencing (CCS) analysis in SMRT Link (v8.0) (https://www.pacb.com/support/software-downloads/) with the following parameters: “--min-passes 1 --min-length 100 --min-rq=0.99”. We first used HiFi reads to create the initial assembly via Hifiasm^20^ software (v0.16.1) with default parameters. Then, the initial assembly was screened against the NCBI nonredundant nucleotide (Nt) database to remove mitochondrial sequences and bacterial contaminants using the BLASTN^71^ tool (v2.10.0).

### Bionano scaffolding

The Bionano data analysis, including data filtering, *de novo* assembly and scaffolding, was performed using the Bionano Solve software suite (v3.7.1, https://bionano.com/software-downloads/). In brief, Bionano raw data were quality controlled with a molecular length of <150 kb, a signal-to-noise ratio (SNR) of <2.75 and a label intensity of >0.8. *De novo* assembly of clean Bionano data was performed to generate consensus maps using the BioNano Solve software suite (v3.7.1). Hybrid scaffolding of the contigs by Hifiasm and Bionano optical maps was used to obtain superscaffolds. To construct the hybrid scaffold maps, the assembled contigs were converted to *in silico* cmap format and then aligned to the Bionano consensus maps using the proprietary alignment tool RefAligner of Bionano (https://bionano.com/software-downloads/). Finally, scaffold sequences were produced according to the above alignments before subsequent Hi-C anchoring.

### Pseudochromosome construction

Quality control was conducted on the Hi-C raw reads by HiC-Pro^72^ (v2.8.1). Clean Hi-C reads were aligned to the scaffolds produced by Bionano using Bowtie2^73^ (v2.3.2) with default parameters. Valid read pairs were used to place the scaffolds onto pseudochromosomes based on their interactions by using LACHESIS^74^ with default parameters. Potential assembly errors were manually checked and adjusted using Juicebox^75^ (v2.18.0). Finally, all the scaffolds were anchored to 27 pseudochromosomes (26 autosomes and the X chromosome), which was consistent with the karyotype results in previous sheep studies^76,77^. We conducted an independent *de novo* T2T assembly of the Y chromosome, which is described in detail below.

### Gap verification and filling

The gaps were verified by aligning HiFi and ONT ultralong reads to pseudochromosomes and manual review of the sequencing coverage in IGV (v2.13)^78^. Some gaps were produced by splitting the conflict sites between the Bionano consensus maps and the initial assembly and were examined manually for potential errors. The gaps were removed by replacement with the original assembled contigs in the initial assembly if the read coverage (≥5) on the original contigs was continuous.

The ≥100-kb ultralong ONT reads within the gaps were searched out based on alignment against the genome assembly via Minimap2^79^ (v2.23) with the parameter “-x map-ont”. Short gaps were filled by extending the overlapping ONT long reads, while the other gaps were further filled by using local assembly based on ONT reads. In brief, the reads that uniquely aligned with the ends of two neighboring contigs connecting the beginning site and ending site of a gap (identity ≥ 95%, coverage ≥ 90%, and QV ≥ 20) were used as anchors, and based on these anchors, the long reads that overlapped themselves (identity < 95% and coverage < 90%) were searched iteratively. Local assembly was then performed based on all the gap-related ONT reads, including the above aligned reads and the reads unmapped to the genome sequences. The *k*-mers (*k* = 23) were generated based on the MGI short reads using Jellyfish^80^ (v2.3.0), and low-frequency *k*-mers of less than the average depth were selected as rare *k*-mers for each ONT read. A string assembly graph was built based on the overlapping ONT reads and their rare *k*-mers using the NextGraph module in NextDenovo^81^ (v2.5.2). The final graph was reached when the longest accumulated length of rare *k*-mers was achieved, and a contig was obtained accordingly to connect the beginning site and ending site of a gap. The gap-filled pseudochromosomes were double checked to ensure the correct gap-free genome assembly with possible manual adjustment, based on the alignments and coverage of all the ONT reads in IGV (v2.13).

### Initial assembly of the Y chromosome

The haplotype-resolved assembly strategy was adopted to assemble the Y chromosome following a modified version of Koren’s method^12^. Using MGI whole-genome shotgun sequencing data, *21*-mer libraries unique to HU-3095 and its parents were constructed using Jellyfish^80^ (v2.3.0). Paternal *21*-mers in HU3095 were identified based on their unique presence in the father but not in the mother. Paternal ultralong ONT reads (1.73 Gb, ∼64.13× coverage) were chosen based on more paternal *21*-mers than maternal ones. The paternal ONT reads that were uniquely aligned to the autosomes by Minimap2 (v2.23) were removed. The remaining ONT reads potentially from the paternal X and Y chromosomes were used to construct an assembly graph of the Y chromosome using the NextGraph module in NextDenovo^81^ (v2.5.2). The assembled graph was manually adjusted for gap filling, scaffolding, and correction, with assistance from the Y contigs using the trio-binning option of Hifiasm (v0.16.1) based on HiFi reads and *31*-mers of the parents of HU3095. To validate the completeness and reliability of the above initial Y chromosome assembly, the Y chromosome (CP128831.1) from *Ramb_v3.0* was included in the collinearity analysis using MUMmer^82^ (v4.0.0), and the length of the Y chromosome assembled here was double checked based on the estimated karyotype length in previous studies^76,77^. Alignment of Bionano consensus maps against the Y chromosome was performed to examine possible assembly errors.

### Telomere filling

HiFi reads containing >10 copies of the telomere-specific repeat sequence “AACCCT/AGGGTT” were retrieved as type I using BLASTN (v2.10.0). The HiFi reads were aligned to the gap-free genome assembly using Minimap2 (v2.26), and those without any hits against the genome were extracted as type II using SAMtools (v1.18)^83^. The type III HiFi reads could be aligned to a 1-Mb interval at the chromosomal ends. The above three types of HiFi reads (types I, II, and III) were used to construct the primary assembly of telomeric regions using Hifiasm (v0.16.1) with default parameters. The contigs were corrected and scaffolded together with the sequences of 1-Mb chromosomal ends using RagTag^84^ (v2.1.0), and the telomeres were placed at the two ends of each chromosome (Supplementary Methods).

### Genome polishing

Genome polishing was performed using a customized pipeline (https://github.com/lly1214/CAU-T2T-Sheep), which included five steps of polishing (Supplementary Methods). HiFi reads were first mapped to the gap-free genome assembly using Minimap2 (v2.26)^79^. The low-quality regions (LQRs) were determined based on the three cutoffs of a mapping quality (MAPQ) score ≤1, clipped reads identified at their two ends, and <3 HiFi-aligned reads, when compared to the high-quality regions (HQRs). These LQRs were polished in the first round with HiFi reads using NextPolish2 (v0.2.0)^85^ and two additional independent rounds with ultralong ONT and MGI reads using NextPolish (v1.4.1)^86^. LQRs and HQRs were merged into one whole genome before the last round of NextPolish2 (v0.2.0) polishing based on HiFi long reads. Finally, a gap-free complete genome assembly (*T2T-sheep1.0*) of all chromosomes, including autosomes and chromosomes X and Y, was constructed for sheep, with the average QV (51.53) higher than that (36.30) for the unpolished genome sequences.

### Haplotype genome assembly

The trio strategy was used to assemble the autosomes of the haplotype-resolved genomes (*T2T-sheep1.0P* and *T2T-sheep1.0M*), based on HiFi reads and parents’ short reads, by using Hifiasm (v0.16.1 r375) with the default parameters. The scaffolding, gap filling and polishing for *T2T-sheep1.0P* and *T2T-sheep1.0M* were performed according to the similar method for *T2T-sheep1.0*. ONT and HiFi reads were trio-binned and determined for paternal and maternal origins based on the *21*-mers from parents’ short reads and the paternal or maternal dominance. The binned ONT and HiFi reads were used for filling gaps and polishing. The haplotype genomes were polished based on binned ONT and HiFi reads for two rounds by using NextPolish2 (v0.2.0), and the NGS data was not involved to avoid the introduction of the other haplotype sequences.

### Assessment and validation of genome assemblies

The *T2T-sheep1.0* genome assembly was validated by multiple methods, including the coverage of reads, collinearity and QV. Depth coverage was calculated in 200-kb windows using Bamdst (https://github.com/shiquan/bamdst) based on the bam files of the HiFi and ONT long reads by Minimap2 (v2.26) and short reads by BWA^87^ (v0.7.17). The genome coverage was plotted using the karyoploteR^88^ package (v1.8.4). The reliability of the *T2T-sheep1.0* assembly was compared with that of the most updated sheep genome reference, *ARS-UI_Ramb_v3.0* (GCF_016772045.2), based on collinearity analysis, which was performed based on alignment using Minimap2 (v2.26) with the parameter “-cx asm10”. The genome synteny between the two assemblies was visualized using paf2doplot (https://github.com/moold/paf2dotplot). A *21*-mer hash table was created from the MGI short reads using the meryl command of Merqury^89^ (v1.3.1). The quality score (QV), switch error, and error *k*-mer frequency of *T2T-sheep1.0* and the haplotypes were calculated accordingly. In addition, we downloaded 26 published chromosome-level ovine genomes (Supplementary Table 5), and compared them with *T2T-sheep1.0* for gap lengths, gap counts, total bases, total bases with unplaced contigs excluded, and total bases with the mitochondrial genomes and unknown bases of gaps excluded.

The completeness of the *T2T-sheep1.0* assembly was assessed using BUSCO^90^ (v4.0.5) based on the mammalia_odb10 database. To evaluate the accuracy of *T2T-sheep1.0*, all the MGI paired-end reads were mapped to the assembly using BWA (v0.7.17). We computed the mapping rate and base accuracy using SAMtools^83^ (v1.18) and BCFtools^83^ (v1.15.1).

### Repeat annotation

Tandem repeats were *de novo* predicted using GMATA^91^ (v2.2) and Tandem Repeats Finder^92^ (TRF, v4.09.1). Transposable elements were *de novo* predicted using MITE-Hunter^93^ (v1.0) and RepeatModeler2^94^ (v2.0.4). The *de novo* repeat libraries were merged with the Repbase^95^ database of repetitive DNA elements. The merged repeat library was then used to perform repeat annotation using RepeatMasker (v4.1.4)^96^ with default parameters. The repeats were masked in the *T2T-sheep1.0* assembly using RepeatMasker, and used for subsequent SDs and coding-gene annotation.

### Segmental duplication (SD) identification

First, the SDs were detected using BISER^97^ (v1.4) with the parameters: “--max-error 20 -- max-edit-error 10 --kmer-size 31”. Afterward, the SDs were filtered following a previously described method for human *T2T-CHM13*^98^. In brief, filtering was based on the following criteria: >90% gap-compressed identity, ≤50% gapped sequence in the alignment, >1 kb of the aligned sequence, and ≤70% satellite sequence as assessed by RepeatMasker. Finally, SDs were plotted using Circos^99^ (v0.69). We counted the number of SDs that overlapped with PURs and genes using local scripts.

### Protein-coding gene annotation

A combination of *de novo* prediction, homolog-based determination, and transcriptome-based identification was used to annotate genes in *T2T-sheep1.0*. For the transcriptome-based approach, RNA-seq data for 28 tissues of Hu sheep (Supplementary Table 7) downloaded from the NCBI were used to assemble the transcripts. In summary, after filtering and quality control, the clean reads were aligned to *T2T-sheep1.0* with STAR^100^ (v2.7.9a), and the transcripts were assembled with Stringtie^101^ (v1.3.4d). Full-length transcripts from Iso-seq were aligned to *T2T-sheep1.0* using Minimap2 (v2.16) with the parameter “-x splice -uf”, and nonredundant transcripts were obtained using the “collapse_isoforms_by_sam.py” command of IsoSeq3 (v3.8.2, https://github.com/PacificBiosciences/IsoSeq). Nonredudant transcripts from RNA-seq and Iso-seq were used to predict gene models via PASA software (v2.5.2)^102^. For the homolog-based approach, homologous proteins from sheep and other mammalian species (e.g., sheep, *O. aries*; argali, *O. ammon*; goat, *Capra hircus*; house mouse, *Mus musculus*; cattle, *Bos taurus*; and human, *Homo sapiens*) were downloaded from the NCBI (Supplementary Table 22), and genes were identified after alignment to the *T2T-sheep1.0* using GeMoMa software^103^. For *de novo* prediction, genes were predicted using the software AUGUSTUS (v3.3.1)^104^ based on the above complete genes. All the above gene models were integrated using EvidenceModeler (v1.1.1)^105^ with default parameters. The genes containing TEs were filtered out using TransposonPSI software (v1.0.0, https://github.com/NBISweden/TransposonPSI). We collected 369 known genes from previous publications and the homologous genes annotated in *Ramb_v3.0*, and manually adjusted gene structures in our gene annotation files in IGV-GSAman (https://gitee.com/CJchen/IGV-sRNA), based on the transcript evidence (RNA-seq and Iso-seq).

### Methylation by PacBio and ONT long reads

Methylated cytosine was examined based on the ONT and HiFi raw data. The HiFi data in BAM format was aligned to the *T2T-sheep1.0* assembly using pbmm2 (v1.13.0) (https://github.com/PacificBiosciences/pbmm2). Subsequently, 5mC methylation probability was generated for the sites using the “aligned_bam_to_cpg_scores” command of pb-CpG-tools (v2.3.1) (https://github.com/PacificBiosciences/pb-CpG-tools). For methylation analysis based on ONT data, Fast5 format files were converted to fastq files using GuPPy^106^ (v6.1.2), and methylated sites were called using Nanopolish^107^ (v0.14.0). We filtered out the methylation sites with a frequency < 50% and calculated the frequency of methylated bases in 10-kb windows using BEDTools^108^ (v2.31.0). Their distribution was plotted with the R package karyoploteR (v1.8.4).

### Identification and validation of centromeric regions

Centromeric regions were first identified by ChIP-seq based on the enrichment of histone binding. The raw ChIP-seq reads were trimmed using fastp (v0.23.1)^109^ with the parameters “-f 5 -F 5 -t 5 -T 5”. Clean ChIP-seq reads were aligned to *T2T-sheep1.0* using Bowtie2 (v2.4.2) with the parameters “–very-sensitive –no-mixed –no-discordant -k 10”. The ChIP-seq peaks corresponding to the centromeric regions were called using MACS3 (v3.0.0b2)^110^, and the average read depth for ChIP enrichment in 10-kb sliding windows was calculated using BEDTools (v2.30.0) and plotted with the karyoploteR package (v1.8.4).

Centromeric regions were further validated based on sequence complexity and identity across the whole genome of *T2T-sheep1.0*. Sequence linguistic complexity and Shannon entropy measures were calculated across the whole chromosomes with NeSSie^111^ in a window size of 1 kb and a step size of 8 bp, where lower values indicate more repetitive sequences. The locations of centromeres with enriched repeats are shown by low entropy values^48^. Sequence similarity within and around the centromeric regions was calculated in a window size of 5 kb, and was used to construct heatmaps in StainedGlass^112^ (v0.5).

### Repeat identification within centromeres

The centromeric regions were estimated based on the above ChIP-seq peaks and enriched alignment of known centromere-specific satellite DNA sequences (KM272303.1). The centromeric sequences on autosomes and the X chromosome were obtained using BEDTools (v2.30.0). Based on the methylation enrichment and the nature of the metacentric centromeres on the Y chromosome, we retrieved a region of ∼5 Mb in the middle of the Y chromosome as the candidate centromeric region. We pooled all the sequences of the candidate centromeric regions and identified novel centromere-specific satellite repeat sequences using Satellite Repeat Finder (SRF)^113^. These satellites were classified into four types according to four minimal repeat units (SatI, 816 bp; SatII, 702 bp; SatIII, 22 bp; and CenY, 2516 bp) based on sequence identity (Supplementary Methods). The abundance of satellite repeats in the centromeric regions was assessed using BLASTN (v2.10.0).

### SV identification based on long-read sequences

To assess the performance of the reference genome for mapping long reads and calling variants, three commonly used SV callers, i.e., Sniffles (v2.0.6, https://github.com/fritzsedlazeck/Sniffles), cuteSV^114^ (v2.0.1) and pbsv (v2.9.0, https://github.com/PacificBiosciences/pbsv), were used to detect SVs. The PacBio HiFi and CLR long-read data of 18 wild and domestic sheep (Supplementary Table 11) were aligned to *T2T-sheep1.0* and *Ramb_v3.0*, respectively, using Minimap2 (v2.26) with the parameters “-x map-pb” for PacBio CLR reads and “-x map-hifi” for PacBio HiFi reads. The sequence depths of the 18 individuals were calculated using the “stat” module of SAMtools (v1.18)^83^. The means and standard deviations of the sequence depths across the 18 individuals were summarized for satellites, genes, and syntenic and nonsyntenic regions, and differences in these indices were assessed when the sequences were mapped against *T2T-sheep1.0* and *Ramb_v3.0* using Mosdepth^115^ (v0.3.4) and 500-bp windows. SVs were called using pbsv and Sniffles with default parameters and using cuteSV with the parameters “--max_cluster_bias_INS 1000 --diff_ratio_merging_INS 0.9 --max_cluster_bias_DEL 1000 -- diff_ratio_merging_DEL 0.5 --genotype”. SVs passing the quality filters suggested by pbsv (flag PASS), cuteSV (flag PASS) and Sniffles (flag PRECISE) were retained for merging that was performed by using SURVIVOR (v1.0.7)^116^ with the parameters “1000 2 1 1 0 50”. Only SVs supported by more than two software tools were kept, and further merged across all 18 sheep samples using SURVIVOR (v1.0.7) with the parameters “1000 1 1 1 0 50”.

### SNP and SV calling based on short-read sequences

To assess the performance of short-read alignment and variant calling using *T2T-sheep1.0* as the reference genome, we collected whole-genome sequencing data from 810 wild and domestic sheep across the world from the NCBI database (Supplementary Table 15). Low-quality bases and reads were removed using Trimmomatic^117^ (v0.39), and the high-quality paired-end reads were aligned to *T2T-sheep1.0* using BWA mem (v0.7.17-r1188) with default parameters. The mapped reads were then converted into bam files and sorted using SAMtools^83^ (v1.16). The sequence depth and mapping statistics were summarized using the “stats” module of SAMtools.

SNPs for individuals were called from the bam files using the module “HaplotypeCaller” and were merged using the modules “GenomicsDBImport” and “GenotypeGVCFs” in GATK^118^ (v4.3). To reduce potential false-positive calls, “VariantFiltration” of GATK was applied to filter SNPs with the following parameters: “QUAL< 30.0 || QD< 2.0 || MQ< 40.0 || FS> 60.0 || SOR> 3.0 || MQRankSum< -12.5 || ReadPosRankSum< -8.0”. We counted homozygous and heterozygous SNPs for each wild species or sheep population using local scripts. All identified SNPs were annotated in specific genes of *T2T-sheep1.*0 using SnpEff^119^ (v5.1d).

Only samples (534 sheep) with a sequencing depth >15× of short reads were selected to call SVs from the bam files using three tools, namely, Delly^120^ (v0.8.7), Manta^121^ (v1.6.0) and Smoove (v0.2.8, https://github.com/brentp/smoove) with default parameters. SVs called based on short reads were filtered according to the criteria of 50 bp ≤ SVs ≤ 1 Mb and support by more than two software tools. SVs were merged across the 497 domestic sheep and 37 wild sheep by using SURVIVOR (v1.0.7) for a 1-kb merging bin and a minimum length of 50 bp. For populational analysis, SNPs and SVs with a MAF > 0.05 and a proportion of missing genotypes < 10% were retrieved using VCFtools^83^ (v0.1.16).

### Genetic diversity and population structure

High-quality SNPs were used to assess nucleotide diversity (π). The π values were calculated in 1-Mb windows using VCFtools^83^ (v0.1.16). We implemented linkage disequilibrium (LD) pruning to remove SNPs in LD using PLINK^122^ (v2.00a3.7) with the parameters: “--indep-pairwise 50 5 0.2”, and independent SNPs were used for population structure analysis. Principal component analysis (PCA) was performed among the 810 wild and domestic sheep individuals with the “smartpca” command of the EIGENSOFT^123^ package (v8.0.0) and default parameters. Population structure was further validated using ADMIXTURE^124^ (v1.3.0). The independent SNPs were used to generate a genetic distance matrix with VCF2Dis (v1.47, https://github.com/BGI-shenzhen/VCF2Dis) software, and a neighbor-joining (NJ) tree was built using TreeBeST^125^ (v1.9.2) and visualized by iTOL^126^ (v6.8.1). The Neighbor-Net graph based on pairwise *F*_ST_ genetic distances by VCFtools (v0.1.16) was created by using SplitsTree (v6.3.27) software^127^.

### Selective sweeps associated with domestication and selection for wool fineness

As described previously^4,36^, we identified selective sweeps associated with domestication and selection for the wool fineness trait through two methods applied to SNPs: the cross-population composite likelihood ratio (XP-CLR) and pairwise π ratios. For domestication, we used *T2T-sheep1.0* as a reference to reanalyze the genome sequences of wild sheep of 16 Asiatic mouflons compared with five landrace populations of different geographic origins, including five Dutch Drenthe Heath, ten East-Asian Hu, ten Central-Asian Altay, one African Djallonké, and one Middle Eastern Karakul sheep. These wild and domestic sheep were investigated previously with the sheep assembly *Oar_v4.0* (GCA_000298735.2) as a reference^36^. Selective sweeps for fleece fineness were scanned with *T2T-sheep1.0* as the reference for pairwise comparisons among hairy, coarse-wool, medium-wool, and fine-wool individuals of wild and domestic sheep, all of which were previously investigated with *Ramb_v1.0* as the reference^4^. XP-CLR values were calculated in 10-kb nonoverlapping windows for the selection of fleece fineness and in 50-kb windows with a step of 25 kb for domestication using XP-CLR^128^ (v1.1.2) software with the following parameters: “-L 0.95 -P --size 50000 --step 25000”.

Additionally, *F*_ST_ values were also calculated based on SVs in the above populations to detect the selective sweeps associated with domestication and selection for wool fineness, using VCFtools (v0.1.16). We selected the SVs with the top 1% of *F*_ST_ values as the candidate selective sweeps, and plotted them using the R package ggplot2^129^ (v2.1.1). Candidate regions with the top 1% of XP-CLR and *F*_ST_ values were considered to have signals of selective sweeps. The SNPs and SVs located in these top 1% regions of *Oar_v4.0* and *Ramb_v.1.0* were converted to those in *T2T-goat1.0* using LiftOver (https://hgdownload.cse.ucsc.edu/admin/exe/linux.x86_64/liftOver).

## Acknowledgements

We thank Xueyan Feng (China Agricultural University) for the help on data analysis. This work was supported by grants from the National Key Research and Development Program-Key Projects (2021YFD1200900, 2022YFF1003402, 2021YFF1000703, 2022YFE0113300, 2021YFD1300904), the National Key Research and Development Young Scientists Project (2022YFD1302000), the National Natural Science Foundation of China (nos. 32320103006, 32272845, 31825024, 31661143014, 31972527, 32061133010 and U21A20246), the Project of Northern Agriculture and Livestock Husbandry Technical Innovation Center, Chinese Academy of Agricultural Sciences (BFGJ2022002), and the Strategic Priority Research Program of Chinese Academy of Sciences (No. XDA24030205), and the Second Tibetan Plateau Scientific Expedition and Research Program (STEP; no. 2019QZKK0501).

## Data Availability

The genome assemblies (*T2T-sheep1.0* and its maternal and parental haploid assemblies *T2T-sheep1.0P* and *T2T-sheep1.0M*) and raw sequencing data generated in this study, including PacBio HiFi data, ultralong ONT data, MGI data, Iso-seq data and ChIP-seq data, can be achieved from the Genome Sequence Archive in National Genomics Data Center (https://ngdc.cncb.ac.cn/) under the BioProject accession number PRJCA024127 and NCBI under the BioProject accession number PRJNA1033229.

## Code availability

Custom scripts and codes used in this study are available at GitHub (https://github.com/lly1214/CAU-T2T-Sheep). Software and parameters used are stated in the Supplementary Methods with more details.

## Author contributions

M.-H.L. conceived the project, and M.-H.L. and S.-G.J. supervised the study. L.-Y.L. and H.W. performed genome assembly and data analysis. L.-M.Z. participated in data analysis and manuscript drafting. W.-M.W. provided part help and financial support for the study. Y.-H.Z. was involved in data analysis. J.-H.H. was involved in interpretation and plotting of the results. L.-Y.L. H.-T.W. and Q.-Y.L. performed the blood collection for HU3095 and its parents. For PacBio sequencing to call SVs, H.-H.E. and L.-Q.Z. collected the blood samples of Tan sheep, and L.-Y.L. and D.-X.M. collected the blood samples of European mouflon. S.-G.J., L.-Y.L., and M.-H.L. wrote the manuscript.

## Supplementary Figures

**Supplementary Fig. 1. HU3095 for T2T assembly and statistics of the HiFi and ONT sequencing data. a,** HU3095 for T2T assembly. **b**, Ultralong ONT read length distribution for HU3095. **c,** HiFi read length distribution for HU3095.

**Supplementary Fig. 2. Genomic features of the chromosomes of *T2T-sheep1.0*.** The coverages of ultralong ONT (Cov. ONT) and PacBio HiFi (Cov. HiFi) long reads are shown in 200-kb windows. Tandem repeat (TE, green), gene (blue), and segmental duplication (SD, purple) density values were calculated in 10-kb windows. MUK, minimum unique *k*-mer length in 100-kb windows. PURs (orange), previously unresolved regions in *T2T-sheep1.0* compared to *Ramb_v3.0*. Error *k*-mer (red), *21*-mer errors. Centromeres are highlighted in dark blue, and telomeres are marked with black triangles.

**Supplementary Fig. 3. Bionano and Hi-C validation of *T2T-sheep1.0*. a,** Bionano alignments are shown for the four selected chromosomes (Chr02, Chr11, Chr16, and ChrX). **b,** Hi-C interaction heatmap showing the reliability of all chromosomes in *T2T-sheep1.0*.

**Supplementary Fig. 4. Telomeric lengths on the chromosomal ends of *T2T-sheep1.0.*** The number of “CCCTAA” repetitive units in telomeric regions was summarized in 1-kb windows at both chromosomal ends in *T2T-sheep1.0*.

**Supplementary Fig. 5. Haplotype-resolved assemblies of paternal *T2T-sheep1.0P* and maternal *T2T-sheep1.0M*. a,** Visualization of the heterozygous regions between *T2T-sheep1.0P* and *T2T-sheep1.0M* using bubbles, according to the GitHub scripts (https://github.com/T2T-CN1/CN1/tree/main/heterozygosity). The parameter of h = 4 (count of SVs per 500 Kb) was set as the threshold for displaying bubbles. The homozygous regions are shown as single paths (grey), and the heterozygous regions are marked as bubbles in blue and red colors. Centromeres are marked as black lines. Uniform whole-genome coverage of binned HiFi and ONT reads is shown for *T2T-sheep1.0P* (**b**) and *T2T-sheep1.0M* (**c**). The abnormal coverage regions (>2*the average depth of whole genome or <0.5*the average depth of whole genome) are indicated by triangles for the potential issues.

**Supplementary Fig. 6. Genomic features of chromosomes of *Ramb_v3.0*.** The labels are the same as those in Supplementary Fig. 2.

**Supplementary Fig. 7. Four gaps in *Ramb_v3.0* that have been filled in *T2T-sheep1.0*. a,** The two small gaps on Chr01 and Chr10 do not contain genes. Gaps in *Ramb_v3.0* are marked in red. Purple lines indicate collinearity between *T2T-sheep1.0* and *Ramb_v3.0*. **b,** The genes annotated in two gaps on Chr23 and Chr05 of *Ramb_v3.0*. The genes in *T2T- sheep1.0* and *Ramb_v3.0* are shown in yellow and green, respectively.

**Supplementary Fig. 8. Comparison among the available sheep genome assemblies.** The gap length (**a**), gap number (**b**), BUSCO (**c**), and length of two known centromeric satellite sequences (SatI and SatII, **d**) compared among the 27 available sheep assemblies (sample details in Supplementary Table 5).

**Supplementary Fig. 9. Inversion errors on chromosomes 9 and X of *T2T-sheep1.0* compared with *Ramb_v3.0*.** The gray lines represent collinear regions, and the orange lines represent inversions. The alignments of PacBio HiFi reads were used to check for the inversion errors of INV195 on chromosome 9 (Chr09) and INV405 and INV406 on chromosome X (ChrX), based on the IGV snapshots. PacBio reads from both Rambouillet sheep (NCBI Biosample ID SAMN17575729 for *Ramb_v3.0*) and HU3095 (i.e., *T2T-sheep1.0* individual) were aligned to *T2T-sheep1.0* well, suggesting the correct assembly on the inversion regions, but the alignment of PacBio reads to *Ramb_v3.0* cannot cover the junction sites of these three inversions. **a,** *Ramb_v3.0* reads were aligned to *T2T-sheep1.0* at the junctions of INV195 on Chr09. **b,** *T2T-sheep1.0* reads were aligned to the two assemblies *T2T-sheep1.0* and *Ramb_v3.0* at the junctions of INV195 on Chr09. **c,** Reads from *T2T-sheep1.0* and *Ramb_v3.0* were aligned to *Ramb_v3.0* and *T2T-sheep1.0* respectively at the junctions of INV405 and INV406 on ChrX.

**Supplementary Fig. 10. Minimum unique *k*-mer length per 100 kb on all the chromosomes of *T2T-sheep1.0* and *Ramb_v3.0*.** Minimum unique *k*-mers (MUKs) were calculated in 100-kb windows for *T2T-sheep1.0* and *Ramb_v3.0*, according to T2T Minimum Unique K-mer Analysis pipeline (https://github.com/msauria/T2T_MUK_Analysis). The more MUK values indicate more repetitive sequences in a 100-kb window.

**Supplementary Fig. 11. Transcriptional expression of genes in previously unresolved regions (PURs) and newly assembled genes of *T2T-sheep1.0*. a,** Newly assembled genes (NAGs) of *T2T-sheep1.0* compared to all sequences of *Ramb_v3.0*. **b**, Genes in PURs in *T2T-sheep1.0* compared to only the chromosomes of *Ramb_v3.0*. **c**, Expression of NAGs in different tissues. **d**, Expression of genes in PURs in different tissues.

**Supplementary Fig. 12. Circos plot for SDs and genes in orthogroups.** From the outer to the inner layer: 28 chromosomes of *T2T-sheep1.0* (**a**), density of genes in orthogroups identified by the OrthoFinder software (**b**), selected genes based on SNPs and SVs associated with domestication (**c**) and wool fineness (**d**) and SDs (**e**). SDs in PURs are highlighted in red, interchromosomal SDs in gray, and intrachromosomal SDs in black.

**Supplementary Fig. 13. Centromeric regions for selected autosomes and ChrX.** Chr03 (**a**), Chr04 (**b**), Chr10 (**c**), and ChrX (**d**) are selected to show the centromeric features. From top to bottom: methylation based on HiFi reads; ChIP-seq for histone H3 variant CENP-A (phospho-CENP-A (Ser7) antibody); Centromeric satellite units of SatI (purple), SatII (blue-green), and SatIII (orange); Repeats of satellite (grey purple), LTR (red), LINE (green), and SINE (orange); the chromosome bar with centromere highlighted in blue; and the sequence identity heatmap (bottom) with the color scale at the left bottom corner in nonoverlapping 5-kb windows.

**Supplementary Fig. 14. Entropy plots for the selected chromosomes of *T2T-sheep1.0*.** Entropy values were calculated across the whole chromosomes (Chr03, Chr04, Chr10, and ChrX) with NeSSie software using a sliding window size of 10 kb with a step of 1 kb.

**Supplementary Fig. 15. Variants in SatI and SatII between *T2T-goat1.0* and *T2T-sheep1.0*.** The aligned sequences of SatI (**a**) and SatII (**b**) between *T2T-goat1.0* and *T2T-sheep1.0* are shown.

**Supplementary Fig. 16. Y-chromosome assembly for different animal species and phylogenetic tree of the *ZFY* gene family on the Y chromosome of *T2T-sheep1.0*. a,** Summarized history of Y-chromosome assemblies in major animals, including pig, human, mouse, donkey, cattle, and sheep. **b,** The nucleotide sequences of *ZFY* genes were used to reconstruct a maximum likelihood (ML) phylogenetic tree, with the *ZFY* gene in *T2T-goat1.0* serving as an outgroup.

**Supplementary Fig. 17. Expression heatmap of all genes on chromosome Y of *T2T-sheep1.0* in 28 tissues.**

**Supplementary Fig. 18. Alignment of long reads from 18 sheep and SV calling. a,** European mouflon and Tan sheep for PacBio sequencing which was performed in this study and the subsequent SV calling. **b,** The mean coverage values and their standard deviation (std) among 18 sheep (X axis) were calculated in 500-bp windows in all genomic regions (all), genes (gene), nonsyntenic regions (non-syn), satellite repeats (satellite), and syntenic regions (syn), in a comparison of *T2T-sheep1.0* and *Ramb_v3.0*. In contrast to the PacBio HiFi reads, the PacBio continuous long reads (CLRs) with more sequencing errors from the three sheep exhibited more abnormal coverage when aligned to both assemblies. **c,** The length distribution for the counts of DELs and INSs based on long reads were compared in a line plot (left) between *T2T-sheep1.0* and *Ramb_v3.0* used as references. The total counts (right) of INSs and DELs are compared between *T2T-sheep1.0* and *Ramb_v3.0* as references. **d,** The counts of DELs and INSs of the different lengths based on long reads were compared between *T2T-sheep1.0* and *Ramb_v3.0* as references, in LINEs, SINEs, LTRs, exon, gene, and non-PURs. DEL is colored in blue for *T2T-sheep1.0* and light blue for *Ramb_v3.0*, and INS is colored in red for *T2T-sheep1.0* and orange for *Ramb_v3.0*.

**Supplementary Fig. 19. SV density data obtained from PacBio data using *T2T-sheep1.0* as a reference.** SV density was calculated in 10-kb windows based on the PacBio data in 18 sheep (Supplementary Table 11). Centromeres and telomeres are shown in yellow and black respectively.

**Supplementary Fig. 20. A homologous deletion inside *TUBE1* gene in the 18 samples.** Comparison of *TUBE1* on *T2T-sheep1.0* and *Ramb_v3.0* (top), with exons in red and blue colors supported by Iso-seq reads. Based on the coverages, a deletion allele was detected in all 18 individuals based on PacBio long reads with *T2T-sheep1.0* as the reference (bottom left), while no deletion was found with *Ramb_v3.0* as the reference (bottom right).

**Supplementary Fig. 21. Improvement of short read alignment in *T2T-sheep1.0* compared to *Ramb_v3.0*.** The 810 samples were divided into 6 geographic domestic sheep populations and wild sheep. Compared with *Ramb_v3.0* (orange), *T2T-sheep1.0* (blue) showed improvement of read alignments, including MQ0 (**a**), aligned properly paired reads (**b**), outward oriented pairs (**c**), aligned reads (**d**) and error rate (**e**).

**Supplementary Fig. 22. Statistics of SNPs based on short reads. a,** The SNPs in 810 samples of all 28 chromosomes assessed against *T2T-sheep1.0* and *Ramb_v1.0*. “Total” and “PASS” indicate the SNPs before and after filtering by GATK program, respectively. **b,** The SNPs in PURs on each chromosome in *T2T-sheep1.0*. Total (**c**), heterozygous (**d**) and homozygous (**e**) SNPs for different geographic domestic and wild sheep populations in a comparison of using *T2T-sheep1.0* (red) and *Ramb_v1.0* (green) as references.

**Supplementary Fig. 23. Average nucleotide diversity (π) of domestic and wild sheep determined using *T2T-sheep1.0* as the reference.**

**Supplementary Fig. 24. Neighbor-Net tree for the populations of domestic and wild sheep.** A Neighbor-Net tree was constructed based on SNPs using *T2T-sheep1.0* as the reference and the *F*_ST_ genetic distances among the domestic and wild sheep. To better visualize the detailed breeds’ names, the branches are magnified around the tree, indicated by dashed lines and arrows. Six superpopulations according to the continents are colored.

**Supplementary Fig. 25. Selected genes associated with domestication and their allele frequencies. a,** Allele frequency differences of SNPs within two *ABCC4* genes (Gene10178 and Gene10446) between Asiatic mouflon (MOU, *Ovis orientalis*) and landrace sheep (*Ovis aries*). The five landrace breeds are Drenthe Heathen (DRS) in Europe, Altay (ALS) in Central Asia, Hu sheep (HUS) in East Asia, Djallonké sheep (DJI) in Africa and Karakul sheep (KAR) in the Middle East. **b,** Validation of π values (π-O. orientalis/π-landrace) and allele frequencies of selected genes (*OAS1*, *BNC1*, *SPAG16*, *CD226* and *FAM20C*) in the PURs. Deletions in the selected genes *ADAMTSL3* (**c**) and *SPAG16* (**d**) are under selection, with allele frequency differences between domestic and wild sheep and their read coverages as viewed in IGV.

**Supplementary Fig. 26. Selected genes associated with the wool fineness trait and their allele frequencies.** The π values (π-hair/π-fine wool) and allele frequencies of the previously reported selected genes (*TP63* and *KRT1*, **a**) and the newly identified genes (*DMXL2*, *TARBP1* and *EPS8*, **b**) in non-PURs in this study are shown. Deletions of selected genes in non-PURs (*DMXL2*, **c**) and PURs (*CA1*, **d**) are validated with read alignments in IGV, and their allele frequencies are shown with differences in fine-, coarse-and medium-wool sheep, compared to hairy sheep.

**Supplementary Fig. 27. XP-CLR values based on SNPs for the four sheep populations with various fleece fiber diameters.** In addition to fine-wool vs. hairy sheep, the other five comparisons of fine-wool vs. medium-wool sheep (**a**), fine-wool vs. coarse-wool sheep (**b**), medium-wool vs. coarse-wool sheep (**c**), medium-wool vs. hairy sheep (**d**), and coarse-wool vs. hairy sheep (**e**) were also used to search for selection signals based on SNPs and the XP-CLR approach. Genes detected in the hairy vs. fine-wool comparison are marked in red, and genes in the PURs are marked in blue. The selection signals in PURs are highlighted with green lines.

**Supplementary Fig. 28. Selection of *IRF2BP2* gene and its selected sites associated with various fleece fiber diameters. a,** The π values (π-hair/π-fine wool) confirms the selection of *IRF2BP2* gene on Chr25. **b,** The allele frequencies of one insertion and eight SNPs are different between hair and fine wool populations, suggesting a fine wool selection. **c,** The six newly identified sites (16307402, 16307498, 16308112, 16308153, 16310330, and 16311100) are located in the promoter region and 5’ upstream regulation region. The insertion site of 16302462 is the one previously reported by Demars et al. 2017^40^, and the two ones of 16303394 and 16305442 are from the study by Lv et al. 2022^4^. Linkage disequilibrium (LD) analysis based on *r*^2^ showed the linkage among these alleles in fine wool population.

**Supplementary Fig. 29. *F*_ST_ values based on SVs for the four sheep populations with various fleece fiber diameters.** The five comparisons of coarse-wool vs. fine-wool, coarse-wool vs. hairy, coarse-wool vs. medium-wool, fine-wool vs. medium-wool, and hairy vs. medium-wool were used to search for selection signals based on SVs and *F*_ST_. Genes detected in the hairy vs. fine-wool comparison are marked in red, and genes in the PURs are marked in blue. The selection signals in the PURs are highlighted with green lines.

## Supplementary Tables

**Supplementary Table 1. Summary of the sequencing data in this study.**

**Supplementary Table 2. Statistics for the *T2T-sheep1.0* assembly.**

**Supplementary Table 3. The 139 gaps in the initial assembly of *T2T-sheep1.0*.**

**Supplementary Table 4. Length of PURs and quality values (QVs) for the chromosomes of *T2T-sheep1.0*, *T2T-sheep1.0P* and *T2T-sheep1.0M*.**

**Supplementary Table 5. Statistics of the ovine genome assemblies downloaded from the NCBI in a comparison to *T2T-sheep1.0*.**

**Supplementary Table 6. Genomic features in the whole genome and PURs of *T2T-sheep1.0*.**

**Supplementary Table 7. RNA-seq samples used for gene annotation and validation.**

**Supplementary Table 8. RNA expression (FPKM) of genes in the centromeric regions.**

**Supplementary Table 9. Number of genes in the top orthogroups identified among four genomes (*T2T-sheep1.0*, Argali, *Ramb_v3.0* and *ARS1*) of closely related species.**

**Supplementary Table 10. Satellite sequences of the sixteen species used for phylogenetic tree reconstruction.**

**Supplementary Table 11. Long reads of 18 sheep samples used for calling structural variants (SVs).**

**Supplementary Table 12. The number of structural variants based on PacBio long reads using *T2T-sheep1.0* and *Ramb_v3.0* as references.**

**Supplementary Table 13. Allele frequency of structural variants based on PacBio long reads using *T2T-sheep1.0* and *Ramb_v3.0* as references.**

**Supplementary Table 14. Homozygous structural variants (SVs) in exons detected among 18 individuals with *T2T-sheep1.0*.**

**Supplementary Table 15. Short reads of 810 wild and domestic sheep used for population genetics analysis.**

**Supplementary Table 16. Counts of SNPs and structural variants (SVs) based on *T2T- sheep1.0* and *Ramb_v1.0*.**

**Supplementary Table 17. Putative selected genomic regions associated with domestication based on SNPs and the top 1% of XP-CLR values.**

**Supplementary Table 18. Selected structural variants (SVs) associated with domestication based on the top 1% of *F_S_*_T_ values.**

**Supplementary Table 19. Genomic regions putatively under selection for the wool fineness trait based on SNPs and the top 1% of XP-CLR values.**

**Supplementary Table 20. Selected structural variants (SVs) for the hairy vs. fine-wool sheep comparison based on the top 1% of *F*_ST_ values.**

**Supplementary Table 21. Structural variants (SVs) within previously unresolved regions (PURs) based on short reads and PacBio long reads.**

**Supplementary Table 22. Genome assemblies for sheep and other mammalian species downloaded from the NCBI for gene annotation.**

